# Cooperative control of a DNA origami force sensor

**DOI:** 10.1101/2023.06.26.546608

**Authors:** Ariel Robbins, Hazen Hildebolt, Michael Neuhoff, Peter Beshay, Jessica O. Winter, Carlos E. Castro, Ralf Bundschuh, Michael G. Poirier

## Abstract

Most biomolecular systems are dependent on a complex interplay of forces. Modern force spectroscopy techniques provide means of interrogating these forces. These techniques, however, are not optimized for studies in constrained or crowded environments as they typically require micron-scale beads in the case of magnetic or optical tweezers, or direct attachment to a cantilever in the case of atomic force microscopy. We implement a nanoscale force-sensing device using a DNA origami which is highly customizable in geometry, functionalization, and mechanical properties. The device, referred to as the NanoDyn, functions as a binary (open or closed) force sensor that undergoes a structural transition under an external force. The transition force is tuned with minor alterations of 1 to 3 DNA oligonucleotides and spans tens of picoNewtons (pN). This actuation of the NanoDyn is reversible and the design parameters strongly influence the efficiency of resetting the initial state, with higher stability devices (≳10 pN) resetting more reliably during repeated force-loading cycles. Finally, we show that the opening force can be adjusted in real time by the addition of a single DNA oligonucleotide. These results establish the NanoDyn as a versatile force sensor and provide fundamental insights into how design parameters modulate mechanical and dynamic properties.

## INTRODUCTION

Biomolecular functions are often driven by inter- and intramolecular forces. Thus, elucidating the forces within and between biomolecular systems provides critical insight into the mechanisms of their functions^1–3^. Molecular force spectroscopy has been a powerful approach for probing the interactions that are responsible for these forces and providing mechanistic insight into function^4–7^. However, current force spectroscopy techniques have limitations such as challenges with force measurements in constrained or crowded environments. For instance, both magnetic and optical tweezers necessitate the use of large (>1 µM) beads, which act as handles for applying forces on nanoscale samples^6–9^. Atomic force microscopy requires the sample be attached to a cantilever tip^4, 6, 10–13^. These methodologies are limited to systems where space is available for the handles, which makes it challenging to implement these approaches within cells^14, 15^ and nanofluidic devices^16, 17^. Here we present the development and investigation of a DNA Origami (DO) nanodevice that has the potential to address these limitations.

DO nanotechnology has significant promise in developing nanodevices for complex functions including drug delivery^18–20^, molecular sensing^21, 22^, and probing single molecule dynamics and interactions^23–28^. More specifically, DO has been established as a useful approach for single molecule force sensing, with demonstration of DO devices applying and detecting both tensile and compressive forces^29–32^. Complex and dynamic 3-dimensional DO nanodevices can perform prescribed functions through controlled actuation, making their use precise and reproducible^30, 33–35^. DO devices are biocompatible, functionalizable, and on the nanometer (nm) size scale, which are key characteristics that position them to function within complex nanoscale environments. In addition, the intrinsic modularity of DO devices allows them to be modified and tuned without the need for redesign of the primary structure^36^.

In this study we focus on a DO force sensor, the NanoDyn (ND), which has been previously shown to be sensitive to compressive depletion forces^30^. Hudoba *et al*. introduced the ND as a sensitive reporter of compressive depletion forces due to local molecular crowding on the order of 100 femtoNewtons (fN) and with a lower limit of force detection of 40 fN. Furthermore, this work demonstrated the viability of this device operating in molecularly crowded environments. Here, we build on that research and demonstrate the utility of the ND not only as a highly sensitive reporter of compressive depletion forces, but also as a robust, dynamic device capable of detecting tensile forces from picoNewtons to tens of picoNewtons (pN) where device design parameters allow tunable control of the force response.

Taking advantage of the modular nature of the ND, we show that an individual single stranded DNA molecule, which we refer to as a zipper strand, can be modified to set the force-sensing capabilities and be incorporated after folding and purifying the ND. This allows for rapid and efficient tuning of the device and eschews the need to fold and purify a separate structure for different force applications. We investigated its response to tensile forces and determined that it can be tuned to be sensitive to a range of forces through the adjustment of 1 to 3 zipper strands. We show that the ND detection force can be adjusted between 5-13 pN by changing a single zipper strand within the device. We then demonstrate that by incorporating multiple zippers in parallel, forces of about 30 pN with the potential of even higher forces can be detected. We find that more stable interactions (opening forces ≳ 10 pN) lead to a higher reclosure probability. Finally, we show that the force-sensing range of the ND can be adjusted in real time by iteratively incorporating DNA zippers *in situ*. This study lays the groundwork for a modular and versatile force probe that has the potential to be used in complex biological systems where traditional force spectroscopy techniques are challenging or impractical to implement.

## RESULTS

### A DNA origami (DO) design for a modular and tunable force sensor

As previously reported^30^, the ND is prepared by scaffolded DNA origami^36–38^ where it consists of two origami bundles in a honeycomb lattice linked by six parallel 116 nucleotide (nt) single strand (ss) connections that we refer to as loops. Each loop is configured as either a “force-sensing” loop or a “hinged” loop with the addition of ssDNA molecule(s) (**Fig. 1a**, **Supplementary Fig. S1**). A force-sensing loop contains a zipper strand DNA oligonucleotide with 3 distinct regions, which allows the ND to transition between open and closed states. (i) The “anchor” region binds to one side of a given loop with 30 complementary bases so that this region remains base paired in both the open and closed states. (ii) The “zipper” region binds the opposite side of the loop such that the two origami bundles are constrained in the closed state. The length of the zipper region is varied between 11 nt and 21 nt to influence the opening force of the ND. (iii) The “linker” region is a 5 nt poly-T sequence linking the anchor and zipper regions, that helps reduce steric clash within the closed state of the ND. The hinged loops contain two separate 46nt DNA molecules referred to as “blocking strands” that form two dsDNA regions that are separated by 14 nt of ssDNA between them and leave a 5 nt ssDNA spacer adjacent to the ND barrels. This reduces the impact of secondary structure and entropic elasticity within the hinge loops on force-sensing. The modularity of the ND allows for the zipper strands to be incorporated after folding and purification of the base structure of the ND in a secondary reheating and slow annealing cycle (see **Methods** for details). This allows a single preparation of the ND base structure to be used for multiple ND force-response configurations. The ND folding, purification and zipper incorporation were verified via Agarose Gel Electrophoresis (AGE) (**Supplementary Fig. S2, S3**) and Transmission Electron Microscopy (TEM) (**Fig. 1a**, **Supplementary Fig. S4**).

**Figure 1:**
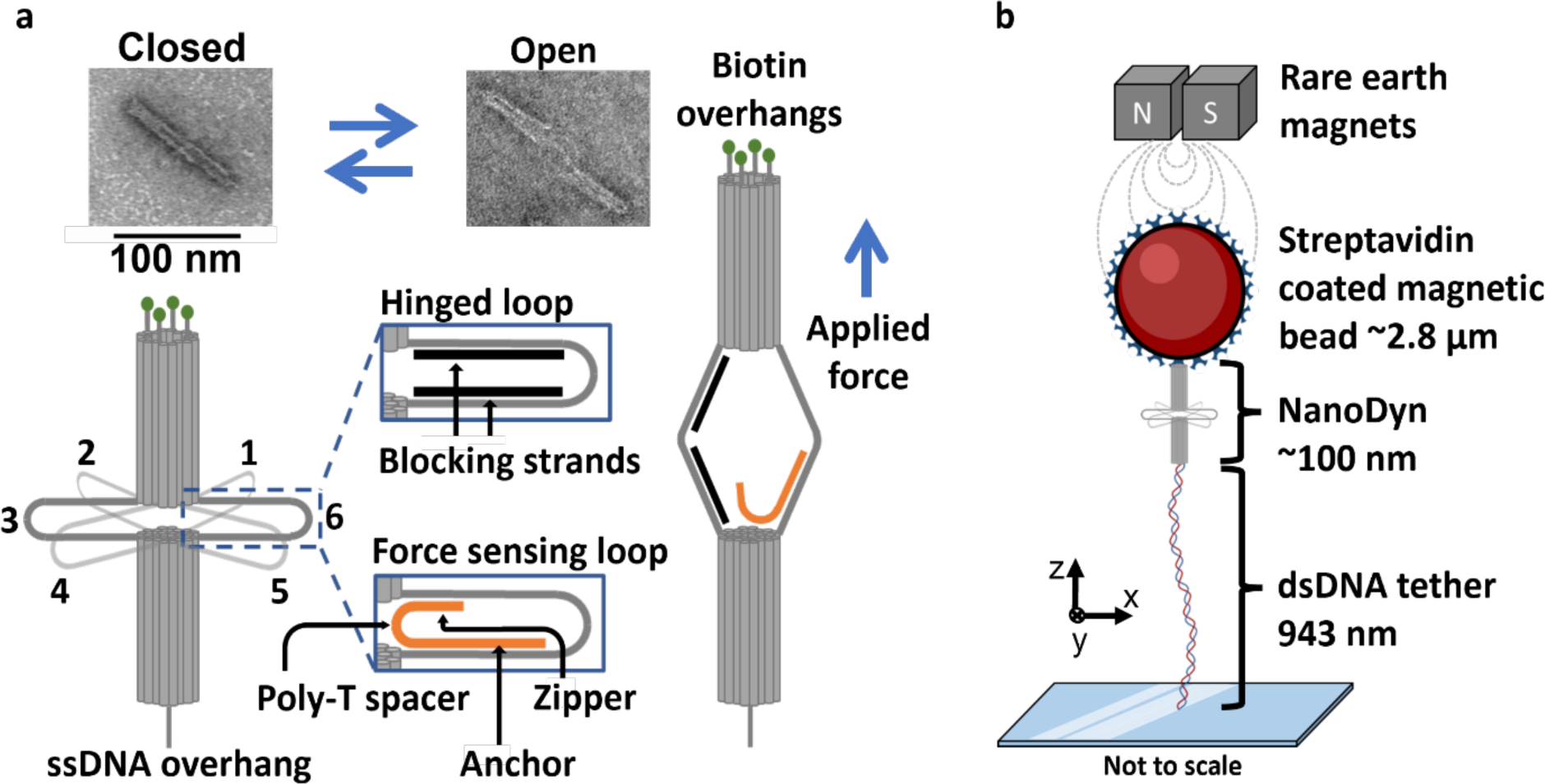
DNA origami NanoDyn schematic and experimental design. (**a**) Schematic drawing of a DNA origami NanoDyn (ND) which consists of 2 honeycomb lattice barrels held together by 6, 116 nt ssDNA crossover strands called loops. Each loop can be folded into a “force-sensing” loop or a “hinged” loop through annealing of a zipper strand or blocking strands, respectively. The force-sensing loop can be in an open or closed state. TEM images provide visualization of these two states. (**b**) The ND is attached to a 2.8 µm superparamagnetic bead through 4 biotin-streptavidin linkages. The opposite end of the ND is annealed to a dsDNA tether, which itself is annealed to an oligonucleotide covalently bonded to a microscope slide via click chemistry. Repeated actuation of the ND is achieved by repeatedly increasing and then decreasing the force using a magnetic tweezers system. A sudden increase in length during the force loading step is indicative of an opening event.

To investigate the force-sensing capabilities of the ND, we used a Magnetic Tweezers (MT) approach to carry out repeated force-extension measurements. Using a lab-built MT on an inverted Olympus IX-70 microscope base, we repeatedly actuated single NDs between the closed and open states by serially increasing and lowering the applied force. Each ND was attached to a 2.8 µm streptavidin coated superparamagnetic bead by 4 biotinylated dsDNA extensions at the top end of the ND (**Fig. 1b**). The opposite end of the ND was anchored to a ∼1 µM dsDNA tether via base pairing between two complementary 30 nt ssDNA overhangs. The opposite end of the tether contained a 60 nt ssDNA overhang that anchored it to a glass slide by annealing to a complementary 60 nt ssDNA oligonucleotide that was covalently bound to the slide surface via click chemistry (see **Methods**). As the force was steadily increased, an abrupt increase in the extension of the ND by tens of nanometers occurred (**Supplementary Fig. S5, Supplementary Table S2**). This indicated an opening event, where the zipper region within the force-sensing loop released from the loop, leading to separation of the two barrel components. This gap size between the open and closed ND agrees with previously reported opening distances^30^. To allow the ND to reclose, the force was decreased to ∼0.5 pN, which allowed the zipper region to rebind within the force-sensing loop and close the ND. This also confirmed that the anchor end remained bound to the loop through any prior opening events. Repeated extension-retraction cycles of multiple NDs resulted in a distribution of opening forces, which allowed identification of the median opening force for a given design.

### A single force-sensing loop can tune the NanoDyn force sensitivity

To investigate the tunability of the force range the ND detects through opening, we first focused on a ND with a single force-sensing loop and varied the zipper region length within this single loop. A schematic of the design is shown in **Fig. 2a**, where a single loop (loop 6) was held closed by a zipper strand. We use the nomenclature ‘L#’ to indicate the loop being referenced, and ‘#nt’ to indicate the length of the zipper region. For example: a 13nt zipper region in loop 6 is referenced as L6-13nt. The remaining 5 loops (L1-L5) were folded as hinged loops (**Fig. 2a**).

**Figure 2:**
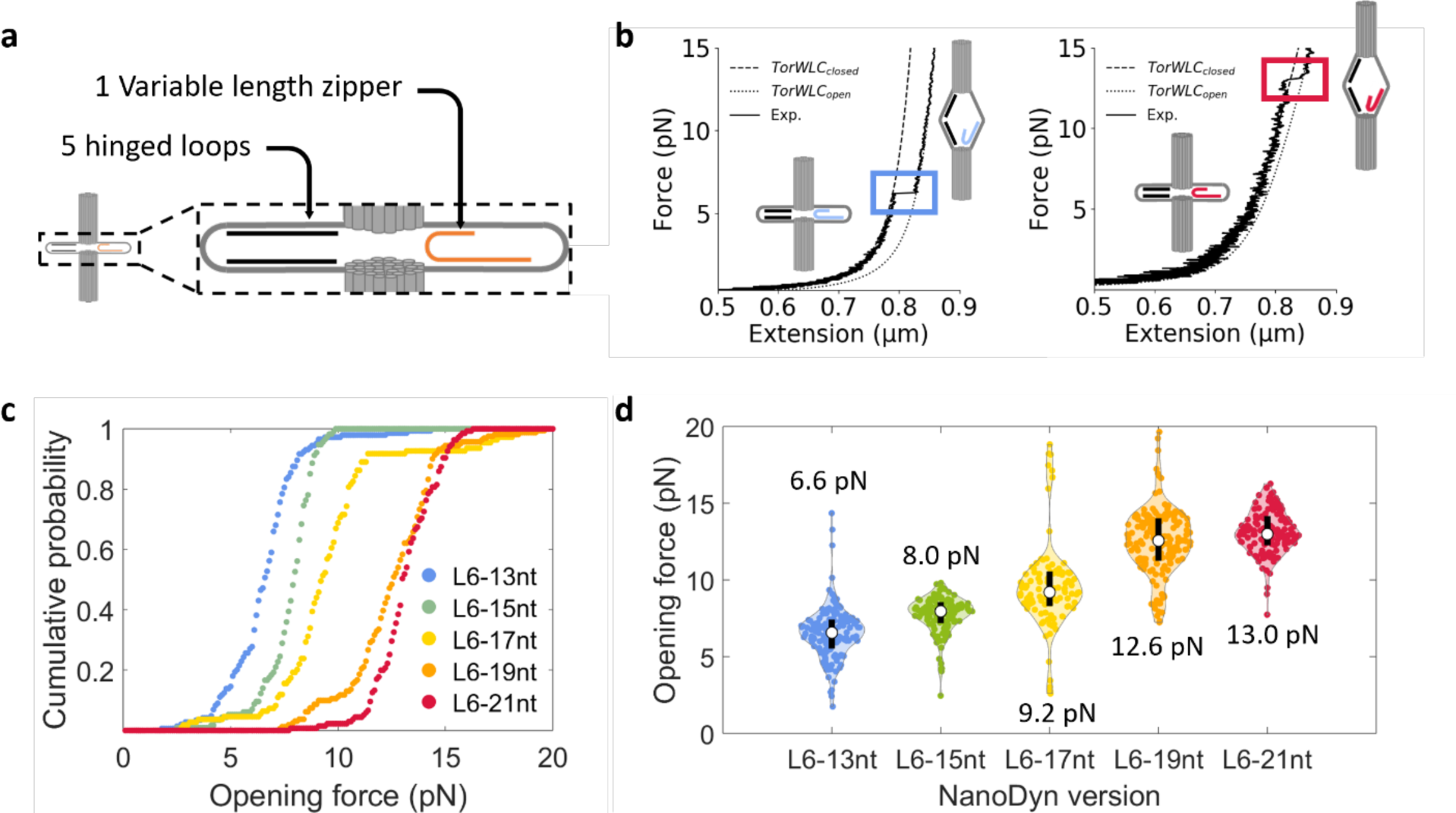
Zipper region length within a single force-sensing loop modulates the opening force. (**a**) Schematic design for ND with one force-sensing loop at L6 and five hinged loops at L1-L5. (**b**) Representative force-extension curves for L6-13nt (blue) and L6-21nt (red). Data was fit to a Torsional spring + worm-like chain model (TorWLC) for both the closed (TorWLC_closed_) and open (TorWLC_open_) states. (**c**) Cumulative probability distribution of the opening forces for multiple pulls across multiple devices. (**d**) Violin plots of the opening force distributions. The white dot is the median opening force value, which is indicated above or below each respective distribution and the black bar indicates the first quartile above and below the median. The median, mean, SD, SEM, and interquartile values for each ND version can be found in **Supplementary Table S1.**

We used the MT to apply repeated force-extension measurements of NDs with a range of zipper region lengths in loop 6. We found that zipper region lengths of less than 13nt in L6 rarely closed. Therefore, we investigated NDs with 5 separate zipper region lengths in L6 of 13nt and longer: L6-13nt (blue), L6-15nt (green), L6-17nt (yellow), L6-19nt (orange), and L6-21nt (red). As the force was increased, the end-to-end extension of the DNA handles with the ND increased continuously with an abrupt extension that was due to the ND opening. Representative force-extension data for L6-13nt and L6-21nt are shown in **Fig. 2b**, while representative data for L6-15nt, L6-17nt, and L6-19nt are shown in **Supplementary Fig. S6**. Because the ND is directly attached to the bead, we develop a worm-like chain plus torsional spring (TorWLC) model to account for the torque applied to the ND as a result of randomized placement of the ND on the bead surface and the alignment of the magnetic moment of the bead with the externally applied magnetic field. (See **Methods** and **Supplementary Methods** for details). The force response of the ND plus DNA handles before the ND opened was fit to the TorWLC_closed_ model (**Supplementary Fig. S7**), while the force response after the ND opened was fit with TorWLC_open_, which included the increased length of the opened ND.

We carried out multiple force-extension measurements with more than 10 molecules of each ND configuration, which resulted in the observation of more than 100 opening events for each configuration (**Supplementary Table S1**). We plotted the cumulative probability (**Fig. 2c**) and determined both the median force (**Fig. 2d**) and the opening distance (**Supplementary Fig. S5, Supplementary Table S2**) of each opening event.

We found that the increase in the zipper region length in steps of two nucleotides from 13 nt to 21 nt correlated with an increase in the median opening force of 6.6 pN, 8.0 pN, 9.2 pN, 12.6 pN, and 13.0 pN, respectively (**Supplementary Table S1**). The standard deviation (SD) for each respective configuration was 1.7 pN, 1.2 pN, 2.8 pN, 2.1 pN, and 1.4 pN (**Supplementary Table S1**). The median, mean, standard error of the mean (SEM), and interquartile values for these opening force and distance measurements are provided in **Supplementary Tables S1** and **S2,** respectively. The observation that most of the opening transitions occurred at or below 15 pN is consistent with previous studies that investigated the forces for unzipping DNA hairpin structures, which show that the force required to unzip DNA converges around 15 to 20 pN^29, 39, 40^ depending on the buffer ionic conditions^41–43^. Overall, these results indicated that length variation of the zipper region within a single loop can finely tune the opening force between 6 and 13 pN, where the lower bound is set by the closure probability and the upper bound is approaching the limit for the unzipping force of a single DNA duplex.

### Parallel force-sensing loops increase the force detect by ND opening

Given the limited range of opening forces the ND can detect with a single force-sensing loop, we investigated the impact of including multiple force-sensing loops arranged in parallel by the ND geometry (**Fig. 3a**). For the second loop, we focused on loop L3 because it is positioned opposite to loop L6, while for the third loop, we used loop L1 that is adjacent to loop L6. We chose zipper region lengths, L1-13nt and L3-11nt so that the melting temperatures (T_m_) were similar (**Supplementary Table S3**) to L6-13nt to help ensure similar opening force medians. For example, L3 has a higher GC percentage, which is why the L3 zipper region was shortened to 11nt. We first prepared the two additional ND versions with a single force-sensing loop at L1-13nt or L3-11nt as controls for studies of NDs with multiple force-sensing loops. We carried out force-extension measurements with the MT (**Fig. 3b** and **Supplementary Fig. S6**), plotted the cumulative probability (**Fig. 3c**), and found that the L1-13nt and L3-11nt NDs opened at respective median forces of 8.5pN and 5.5pN (**Fig. 3d**) with respective SDs of 2.1 pN and 1.2 pN (**Supplementary Table. S1**). The opening force statistics are comparable to the opening forces measured for the L6-13nt ND.

**Figure 3:**
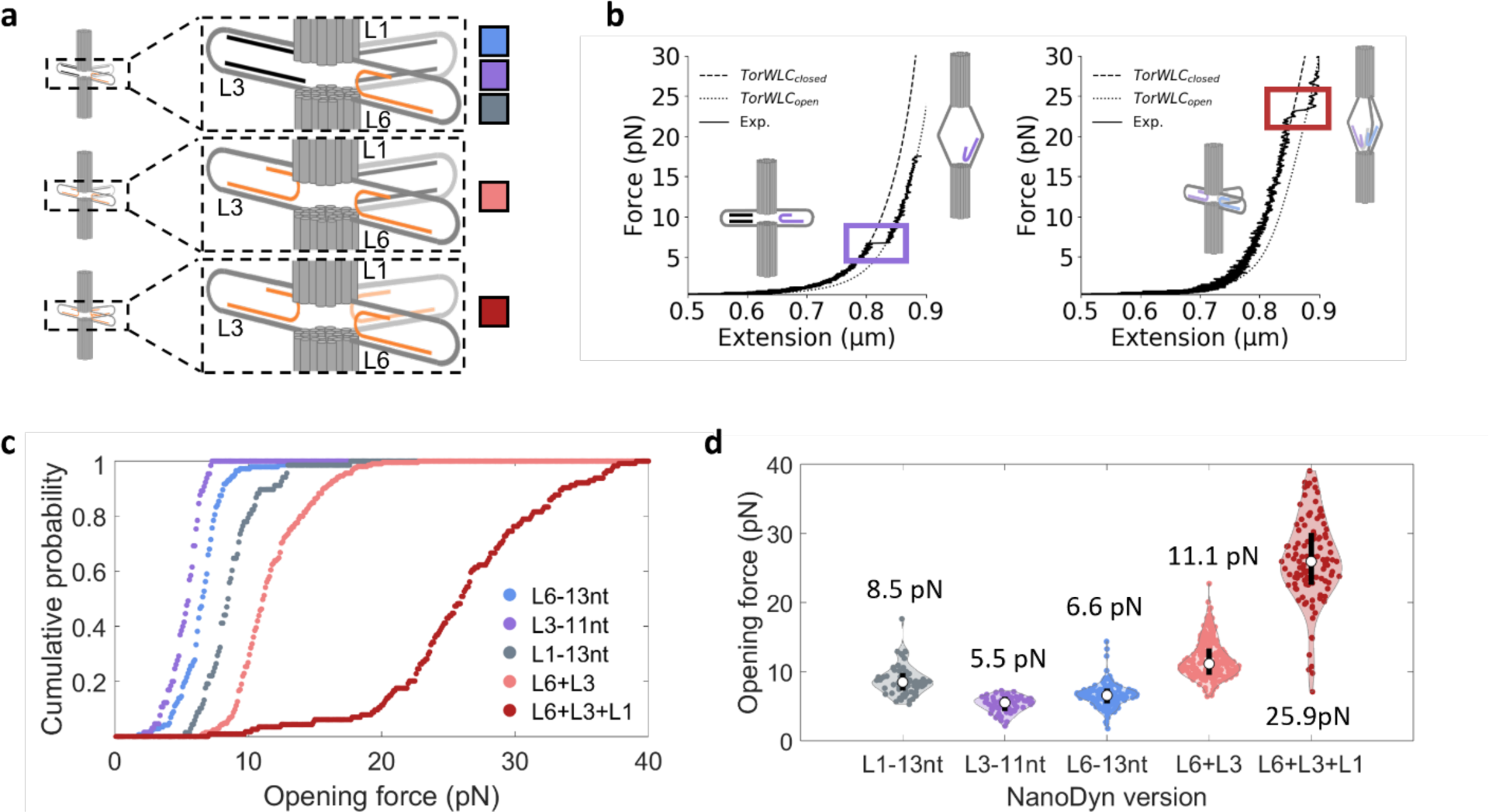
Multiple force-sensing loops increases the ND opening force. (**a**) Schematic design for force-sensing loops with (i) one force-sensing loop at loop 6 (blue), 3 (purple), and 1 (grey); (ii) two force-sensing loops at loops 6 and 3 (pink); and (iii) three force-sensing loops at loops 6, 3, and 1 (dark red). (**b**) Representative force-extension curves for L3-11nt (purple) and L6-13nt + L3-11nt + L1-13nt (dark red - abbreviated L6+L3+L1). Data was fit to the TorWLC model. (**c**) Cumulative probability distributions of the opening forces for the ND devices. (**d**) Violin plots of the opening force distribution for each ND device. The white dot indicates the median opening force, which is shown above or below each respective distribution and the black bar indicates the first quartile above and below the median. The median, mean, SD, SEM, and interquartile values for each ND version can be found in **Supplementary Table S1.**

We then prepared a ND with both L6-13nt and L3-11nt, which we refer to as L6+L3, and investigated the force required to open this two-zipper ND (**Fig. 3c-d**, **Supplementary Fig. S6, Supplementary Table S1**). We carried out force measurements on 23 ND devices and detected a total of 205 opening events (**Supplementary Table S1**). We observed a single step with an opening distance that was nearly identical to the opening of the ND with either L6-13nt or L3-11nt (**Supplementary Fig. S5, Supplementary Table S2**). This is likely because once the first loop opened, the force was significantly above the force that the second closed loop can support, so it opened faster than can be detected by the MT instrument.

For each individual device, there was the possibility that only one of the two zipper strands was incorporated, which would result in a ND that contained only one force-sensing loop. To address this possibility, we verified that each two-zipper ND contained both zipper strands by using log-likelihood analysis (see **Methods** for details). Briefly, by using the two force distributions of a ND with 1 force-sensing loop as the reference model, we determined if any of the force distributions of an individual ND fit better to a one-zipper distribution. Any such devices (5 out of 28 total molecules) were not included in the two-zipper ND analysis. This implies that 82% of the NDs contained 2 force-sensing loops. A histogram comparing the pre and post log-likelihood analysis data is shown in **Supplementary Fig. S8**.

After removal of the devices with a single force-sensing loop, we plotted the cumulative probability of the two-zipper ND opening as a function of force (**Fig. 3c**). We found that the device opened with a single step at a median force of 11.1pN with a SD of 2.8 pN (**Fig. 3d** and **Supplementary Table S1**). For comparison, we plotted the normalized sum of the cumulative probabilities of the ND’s with the single force-sensing loops with either L6-13nt or L3-11nt (**Supplementary Fig. S9, Methods**). We found that this inferred cumulative probability was within 9% of the two-zipper ND that contained both L6-13nt and L3-11nt. Overall, these results indicate that using the ND geometry to orient two force-sensing loops in parallel results in an additive increase in the opening force of the ND, which significantly expands the force-sensing range of the ND. This is somewhat different from a previous study^29^ that used a DO to orient DNA hairpins in parallel. They found that the force to open the hairpins increased but that the increase in force was less than additive. This suggests the ND allows for a more even distribution of forces between zippers.

To build off the results from combining two force-sensing loops in the ND, we included a third zipper strand to introduce a third force-sensing loop, L1-13nt. We carried out force-extension measurements of the three-zipper ND (**Fig. 3b**), which we refer to as L6+L3+L1. We observed a single opening step, as was observed with L6+L3. This is again consistent with the idea that the three force-sensing loops in parallel support a high enough force that once one of the zipper regions releases, the force on the two remaining loops is high enough so they open faster than the time resolution of the MT measurement.

We used log-likelihood analysis (see **Methods** for details) to verify if a L6+L3+L1 ND contained three force-sensing loops, as was done with the L6+L3 ND. We compared the force opening distributions of individual L6+L3+L1 ND to the distributions of each ND with a single force-sensing loop, and the corrected distribution of the L6+L3 ND. We found that 3 out of the 14 devices studied had a distribution that more likely contained two force-sensing loops. Those 3 devices were therefore removed from further analysis. Hence, 11 out of 14 L6+L3+L1 NDs, or 79%, contained all three force-sensing loops. A histogram comparing the pre and post log-likelihood analysis data is shown in **Supplementary Fig. S8**. There remains a small fraction of opening events that appear to align better to the L6+L3 distribution than the L6+L3+L1 distribution. This is likely because when ND reclosed, not all the force-sensing loops reclosed each time. Importantly, this was a small fraction of the total opening events.

After correcting for NDs that contained less than three force-sensing loops, we determined the cumulative probability for the L6+L3+L1 ND to open (**Fig. 3c**) and found that the median opening force was increased to 25.9 pN with a SD of 6.1 pN. We compared these results to the opening force distribution that was inferred from summing the cumulative probabilities of each of the three individual zippers (**Supplementary Fig. S9**), which implied a median opening force of 15.4 pN with a standard deviation of 4.3 pN. This inferred median force was about 40% lower than the measured value, suggesting that additional interactions were introduced into the device by the addition of the third force-sensing loop in L1. Interestingly, the median step size for L6+L3+L1 ND (38.8 nm) was significantly larger (**Supplementary Fig. S5, Supplementary Table S2**) than all the other ND versions we studied including the L6+L3 ND (31.3 nm). Since the length of the ND in the open state will be similar across all configurations, this increase in step size of the L6+L3+L1 ND was likely due to its length in the closed state being shorter than the other ND versions including the L6+L3 ND. This could be due to base stacking interactions between the two barrels of the ND that is known to occur between DNA origami devices^27, 44, 45^ even though the end connections of adjacent dsDNA helices contain ssDNA loops that suppress base stacking interactions. Furthermore, we saw a slight correlation between the opening force and opening distance for L6+L3 and L6+L3+L1 that was not present in the ND’s with only one force-sensing loop (**Supplementary Fig S10**). This is consistent with the idea that additional interactions such as base stacking shorten the length of the closed ND and result in a higher opening force. Overall, these results demonstrate the versatility of the ND, where integrating multiple force-sensing loops in parallel allows the ND to detect forces higher than the inherent limit of single dsDNA unzipping forces, expanding the versatility of this nanoscale device for force-sensing applications.

### Increased interaction stability improves the closure efficiency

For the ND to function as a reversible force sensor that measures repeated application of an external force, it needs to reclose efficiently following release of the applied force. To investigate this, we determined the fraction of times each ND closed (and subsequently reopened) following a reduction in the applied force to 0.5 pN (**Fig. 4a**). We typically did not directly observe a closing event as the force was reduced. This is likely because the closing events usually occurred at forces where the bead height fluctuations were comparable to the closing step size of ∼30 nm, making the closing step difficult to detect (**Supplementary Fig. S11**). So instead, we determined the fraction of force-extension experiments with a subsequent opening event after the force was reduced to 0.5 pN. We found that the opening efficiency of NDs with one force-sensing loop improved as the zipper region length (and force to open) increased (**Fig. 4** and **Supplementary Fig. S12**). For 11 nt and 13 nt zipper regions, the closing efficiency was less than 50% (**Fig. 4, Supplementary Table S4**). An increase in the zipper region length in L6 to 15 nt, 17 nt, 19 nt, and 21 nt resulted in closing efficiencies of 57%, 61%, 99%, and 82%, respectively (**Supplementary Table S4**). This generally implies that increasing the zipper length improved closing efficiency. To rule out the possibility of zipper strand loss over many force-extension cycles of the ND contributed to a lower closing efficiency, we plotted the fraction of devices that re-opened as a function of the cycle number (**Supplementary Fig. S13**). We found that over 8 force-extension cycles, a significant drop in the fraction of devices that re-opened was not observed, indicating that it is unlikely that zipper strands were lost over the course of the experiment, again confirming that the anchor region effectively stayed bound through successive opening events. In the case of L6-21nt, the closing efficiency decreased relative to L6-19nt. One possibility is that the 5-prime end of the zipper region in L6-21 starts with several A’s that could transiently bind to the poly-T region of the zipper strand and interfere with proper rebinding. We determined the opening fraction of L6+L3 and L6+L3+L1 NDs to assess the impact of multiple force-sensing loops (**Fig. 4**). Interestingly, we found that including multiple force-sensing loops reliably increased the closing efficiency to 91% and 94% for the L6+L3 and L6+L3+L1 NDs, respectively. These results indicate that increased stability by either an increase in zipper region length, or inclusion of multiple force-sensing loops improves closing efficiency.

**Figure 4:**
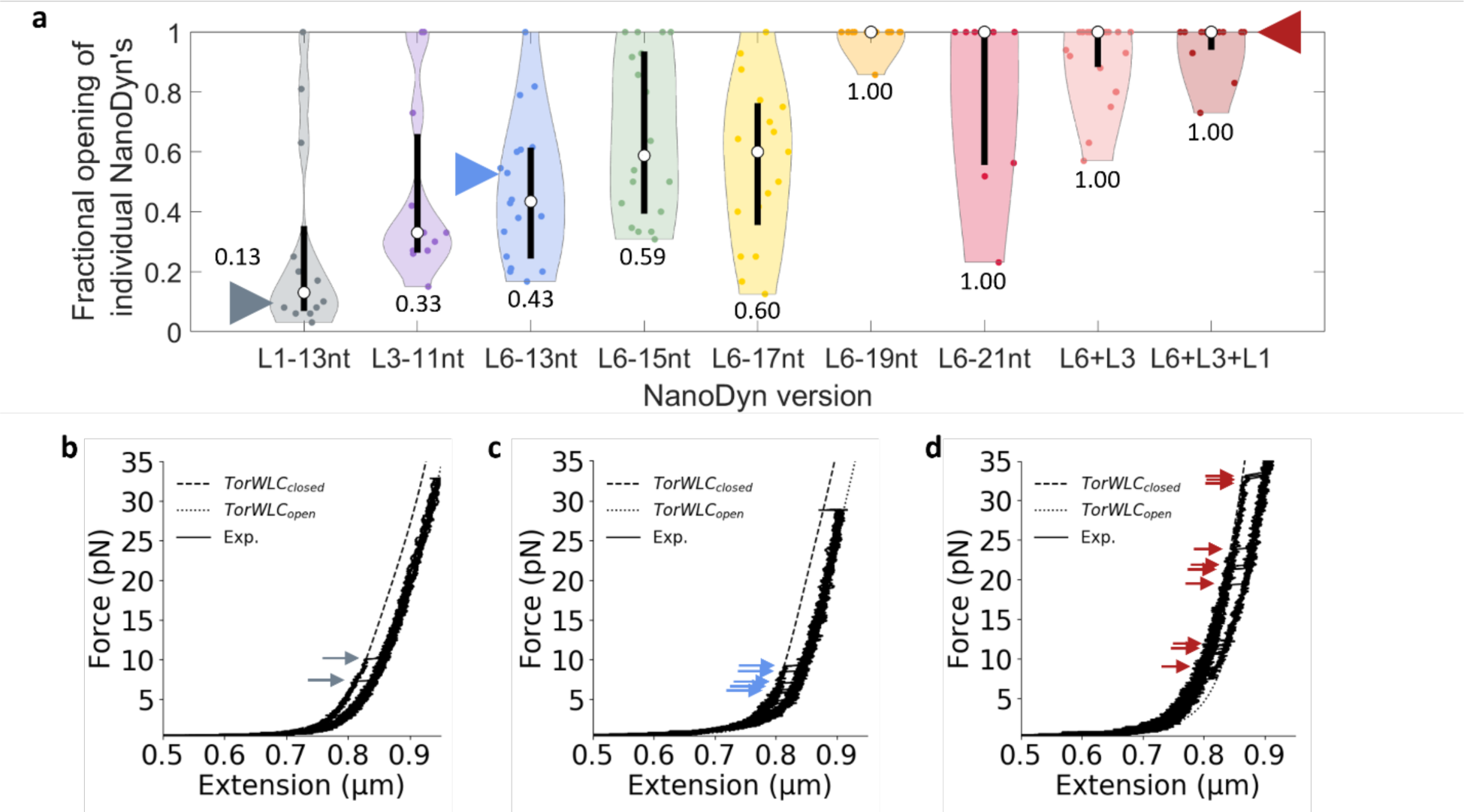
Effect of zipper region length and multiple force-sensing loops on ND closure and subsequent re-opening. (**a**) Violin plots of the fraction of extensions with an opening event for each individual ND. The triangles indicate the position in the distribution of the devices shown in (**b-d**). The white dot indicates the median opening fraction, which is indicated above or below each respective distribution. The black bar indicates the first quartile above and below the median. The median, mean, SD, SEM, and interquartile values for each ND version can be found in **Supplementary Table S4.** (**b-d**) Examples of repeated force-extension curves of single (**b**) L1-13nt, (**c**) L6-13nt, and (**d**) L6+L3+L1 ND devices. Each plot contains 10 consecutive extensions with arrows indicating opening events. The corresponding retraction cycles can be found in **Supplementary Fig. S11**.

### Real-time incorporation of DNA zippers into a single ND to actively control force-sensing

We have shown the force that the ND detects relies on the type and number of force-sensing loops it contains. The zipper strands for the preceding experiments were incorporated after folding of the base structure of the device, which allowed the opening force to be defined through the addition of just a few DNA oligonucleotides prior to use in an experiment. To investigate the potential for real-time modulation of the opening force, we chose an ND design that allowed for successive integration of multiple zipper strands; one to define the initial opening force and another to adjust the opening force of the same single device (**Fig. 5a**).

**Figure 5:**
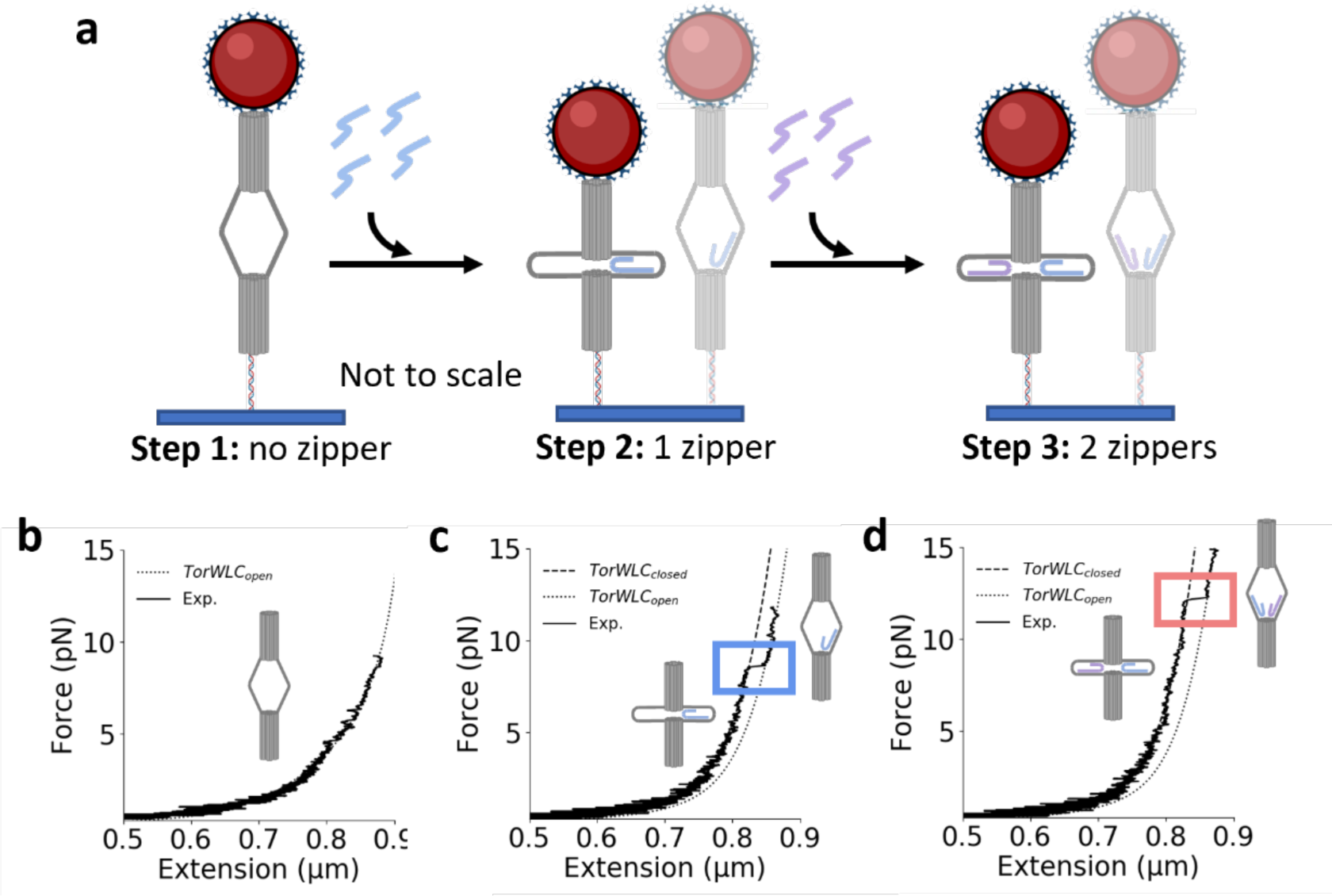
Iterative introduction of multiple force-sensing loops into a single ND. (**a**) Diagram of the iterative introduction of one and then two force-sensing loops into the ND. (**b**) Force-extension of the ND prior to incorporation of a force-sensing loop never results in an opening event. (**c**) Following incubation with a single zipper strand that forms the L6-13nt force-sensing loop, a well-defined step is observed upon force-extension. (**d**) Following incubation with a second zipper strand that forms the L3-11nt force-sensing loop, an additional increase in the opening force is observed.

To demonstrate this approach, we focused on the L6-13nt and L3-11nt zippers, which had median opening forces of 6.6 pN (SD 1.7 pN) and 5.5 pN (SD 1.2 pN), respectively. Starting with an unconstrained ND (omitting blocking strands in L6 and L3), we found that force-extensions never resulted in well-defined opening events, as expected (**Fig. 5b**). We then introduced 800 nM of the L6-13nt zipper strand into the flow chamber and incubated for 20 minutes with a 1 pN applied force on the ND (see **Methods** for details). After flowing out the excess unbound zipper strands, force-extension measurements revealed a single well-defined step in the expected force range (**Fig. 5c**). The opening force was within the force distribution of the L6-13nt ND presented above. To adjust the force-sensing range, we then incubated the ND with a second zipper strand, L3-11nt, 800 nM for 20 minutes and then flowed out any excess zipper strand. Force-extension experiments revealed that the addition of this second zipper strand resulted in an increase in the opening forces to ∼9-13 pN, which is consistent with previous 2-zipper ND measurements (**Fig. 5d**). We found that the first and second zipper strands incorporated with an efficiency of 25% and 23%, respectively. Future work beyond the scope of this study will be necessary to achieve more efficient incorporation of multiple force-sensing loops. However, as a proof-of-concept, these results reveal an important potential avenue for the ND, which is the ability to incorporate zipper strands into the ND *in situ*, allowing for the force-sensing to be initiated and then adjusted within a single device during an experiment.

## DISCUSSION

In this work, we demonstrated that the DNA origami ND can function as a modular nanodevice that can be tuned to detect a range of tensile forces. The modular design allowed the base-structure to be folded, purified, and stored without force-sensing loops. Then, immediately before use, the force detection range was customized by incorporating one or more zipper stands. We demonstrated that varying the zipper region length within a single force-sensing loop modulated the opening force over a range of 3-fold up to a maximum unzipping force of about 15 pN^39, 40^. We then showed that including multiple force-sensing loops in parallel enabled a wider range of opening forces up to 26 pN with only 3 relatively weak zippers, which exceeds the inherent unzipping force of 15 pN. We found that the interaction stability affected the reversibility of ND, with median opening forces below 10 pN not reclosing reliably, indicating that they will not function well for the sensing of repeated force cycling. However, devices with a median opening force above 10 pN repeatedly reclosed, which confirms their utility in detecting repeated force applications. Finally, we showed that single DNA zipper strands can be iteratively incorporated into the ND during a force measurement. This opens the possibility for tuning the force-sensing range of single NDs in real time during a measurement.

This work expands the utility of using nanoscale force sensors as a complementary approach to existing force spectroscopy techniques. The ND has the potential to probe a wide range of forces in constrained environments where it can be difficult to implement other force spectroscopy techniques^4, 6–12^. We previously showed that the ND can operate in crowded environments and detect compressive depletion force in the range of 0.05 to 1 pN^30^. Here in this work, we demonstrated the same ND base structure can also be used to detect tensile forces from 6 to at least 26 pN. The ability for the ND to operate in different modes for detecting both compressive and tensile forces with order of magnitude different force ranges indicates its high versatility for a DNA origami device^23, 25, 26, 29–31, 46, 47^. In comparison to the previous study presented in Dutta *et al.*^29^, our results indicate the ND provides a wider dynamic range of force sensing. This is consistent with the idea that integrating the force-sensing loops between the two-barrel structures allows for a more balanced distribution of the force on these force-sensing loops. There is the potential for further versatility of the ND since up to six force-sensing loops could be included in the ND, each with independent nucleotide sequences to which DNA zippers can be incorporated independently and reproducibly. Assuming an opening force of 10 pN of force per force-sensing loop, using six force-sensing loops should result in an opening force of more than 60pN of force. These large forces do occur in biological systems including the forces on phage genomes during viral packaging^48, 49^ and the forces on mitotic chromosomes during mitosis^50^. However, measurements of these high forces will require covalent attachments or multiple non-covalent attachments to prevent failure of the attachment before device opening and force detection^13, 51^.

The overall length of the ND at 100 nm in length is advantageous for constrained environments. However, for experiments requiring smaller devices, the overall length of the ND could be reduced by designing shorter barrels, while retaining the loop regions. The shortened length could be accomplished with the same DNA scaffold by increasing the width, or with a shorter DNA scaffold. In addition to our current method of monitoring relative length change with magnetic tweezers, a fluorophore pair that undergoes Förster Resonance Energy Transfer (FRET) can be incorporated into the ND with 2 fluorophore labeled oligos, as reported in Hudoba *et al*.^30^, where high FRET reports a closed state and low FRET reports the open state. This will allow detection of a force range in environments where attaching a force handle is not possible.

In the broader context of applications for the ND, there is significant potential for investigating cell-cell and cell-surface interactions based on previous studies. ssDNA hairpins^52, 53^ and DNA origami platforms^29^ have been successfully implemented to investigate intercellular forces. The ND could be used similarly where the modularity of the ND could complement these previously published elegant studies by enabling a wider range of cell adhesion force-sensing. Furthermore, iterative zipper incorporation would allow the force sensor to be tuned in conjunction with changes in the extracellular environment that cause the cells to adapt by changing their cell-surface interactions.

Future studies will be needed to investigate these potential applications of this versatile nanoscale device.

## METHODS

### Preparation of the DNA origami NanoDyn (ND)

The ND design, as previously described in Hudoba *et al*.^30^, consists of 2 DNA bundles connected by 6 crossover strands. One bundle contains 23 dsDNA helices bundled in a honeycomb pattern and has a 30 base ssDNA protruding from the outer end to facilitate attachment to a dsDNA tether. The second bundle contains 17 dsDNA helices arranged in the same fashion as the first bundle but with the central 6 helices omitted. This bundle has 4 biotinylated dsDNA overhangs arranged around the periphery of the outside end (opposite the tethering end) which facilitates attachment to a streptavidin labeled bead. The overall dimension of the ND is ∼100 nm x 15 nm as measured by Transmission Electron Microscopy (TEM). The caDNAno design for this structure is provided in **Supplementary Fig. S1**.

The backbone of the ND consists of an 8064 base scaffold from a modified M13mp18 bacteriophage sequence produced in Castro lab^36^. ∼170 oligonucleotide staples (Integrated DNA Technologies) were combined with the scaffold to fold each version of the device. The exact number of staples is dependent on the version of ND since the number of loops into which zippers are later incorporated will also affect the blocking strands that will be included in the main folding of the ND. Oligo sequences for the base structure can be found in **Supplementary Table S5**, specific zipper sequences in **Supplementary Table S3**, and loop sequences (scaffold crossover points) in **Supplementary Table S6**. The folding was carried out as described in previous folding protocols^36^. In brief, 100 nM scaffold is combined with 10x staple strands in a folding buffer (5 mM Tris, 5 mM NaCl, 1 mM EDTA, 18 mM MgCl_2_, pH 8). The reaction is then subjected to a thermal annealing ramp starting at 65 °C to disrupt non-specific base pairing interactions and then slowly cooled in incremented temperature steps until 4 °C. The detailed thermal ramp is contained in **Supplementary Table S7**. The folding was confirmed by agarose gel electrophoresis (AGE) (**Supplementary Fig. S2**) as previously described^36^. The resulting samples were polyethylene glycol (PEG) purified to remove excess staple strands (**Supplementary Fig. S2**). The PEG purification is based on existing protocols^54^. Briefly, the folded ND sample was combined at a 1:1 volumetric ratio with 15% PEG MW 8000 (Sigma Aldrich) suspended in folding buffer with added 500 mM NaCl. The sample was then pelleted by centrifugation at 16,000 g for 30 minutes at room temperature. The supernatant was removed and the pellet resuspended in folding buffer with 10% glycerol, aliquoted, flash frozen, and stored at – 80 °C^55^.

After folding the base structure of the ND, individual zippers are subsequently incorporated by incubating 4 nM ND with a 10x concentration of the desired zipper(s) for 1 hour at 45 °C then cooling by 1 °C/minute until it reaches 10 °C. This reaction takes place in the same folding buffer described above. In the initial folding, it is recommended to omit blocking strands (strands meant to make the ss scaffold loops double stranded to block unwanted non-specific interactions) only from loops that are intended to be used in subsequent zipper incorporation. The zipper incorporation was confirmed by AGE (**Supplementary Fig. S3**) and TEM (**Supplementary Fig. S4**).

### Preparation of a dsDNA tether handle

Tethers were prepared by restriction enzyme digestion of pUC19 with BsaI (New England Biolabs: R0535) in 1x CutSmart buffer. 1-3 units of enzyme per 1 µg of DNA is usually sufficient to digest the plasmid without over-digestion. The digestion takes place at 37 °C for 1 hour followed by a heat shock at 65 °C for 20 minutes to inactivate the enzyme. The digestion is verified by AGE (**Supplementary Fig. S14**). The gel conditions are as follows and are used for all gels in the tether preparation: 0.7 % agarose gel run in 0.5 xTAE at 225 V and post stained with ethidium bromide for UV visualization. The linearized plasmid was subsequently ligated to oligo pairs to create long ssDNA overhangs (Sigma-Aldrich) on either end. One DNA end has a 3’ 30 nt ss overhang to facilitate attachment to the NanoDyn and the other end has a 3’ 60 base ss overhang to attach to an oligonucleotide covalently bound to the microscope slide. Prior to ligation, the 5’ end of each oligonucleotide to be ligated was phosphorylated using T4 Polynucleotide Kinase (T4PNK) (New England Biolabs: M0201) at 1 U/25 pmol ends incubated at 37 °C for 90 minutes followed by a 65 °C heat shock for 20 minutes. This reaction was performed in 1x T4 ligase buffer rather than T4PNK buffer since they would be subsequently ligated to the linearized plasmid. Each of the 2 oligo pairs were separately annealed at an equal molar ratio at room temperature for 15 minutes in 0.5x TE with 50 mM NaCl (Sigma-Aldrich). The 2 pairs of ends were then ligated simultaneously to the linearized plasmid using T4 DNA Ligase (New England Biolabs:M0202) at 2 U/pmol DNA ends in 1x T4 ligase buffer (provided with enzyme). 100-fold excess ends were used during ligation to prevent recyclization or oligomerization of the linearized plasmid. Ligation was verified by AGE (**Supplementary Fig. S14**). Following the ligation, a phenol chloroform extraction was performed to remove bovine serum albumin (BSA) contained within the digestion reaction buffer. The tether was purified away from excess ends via high pressure liquid chromatography (HPLC) using a Gen-Pak column (Waters: WAT015490). The desired sample fractions were identified by AGE. Those fractions were combined, and the final tether sample was then buffer exchanged and concentrated into 0.5x TE using a 30 kDa centrifugal filter (Millipore-Amicon Ultra).

### Preparation of single molecule experiment slides

The flow cell preparation is an adaptation of the methods used in Luo *et al.*^56^ and Chandradoss *et al.*^57^. First, holes are sandblasted into either end of a coverslip creating an entry and exit port for the sample. This ‘top’ coverslip is then thoroughly rinsed with Milli-Q water to remove excess sand. The top and an equal number of ‘bottom’ coverslips are then sonicated in isopropyl alcohol for 20 minutes. Both sets of coverslips are then thoroughly rinsed in Milli-Q water. The bottom coverslips are dried in an 80 °C oven while the top coverslips are submerged in a 1% Hellmanex III solution (Sigma Z805939), brought to a boil in the microwave, sonicated for 20 minutes, thoroughly rinsed in Milli-Q water, then placed in the 80 °C oven to dry. When the bottom coverslips are dry, they are plasma cleaned (Electron Microscopy Sciences: K100X) for 4 minutes at 25 mA for surface activation. Immediately following the plasma cleaning, the bottom coverslips are incubated in a 3/100 mixture of 3-aminopropyltriethoxysilane (MP Biomedicals: 02154766) and acetone (Sigma 650501) for 30 minutes in a nitrogen gas filled desiccator. Following silanization, the coverslips are sonicated in fresh acetone for 5 minutes, thoroughly rinsed with Milli-Q water, and placed in the 80 °C oven to dry. Once dry, the top and bottom coverslips are sandwiched around parafilm with a channel cut into it. The sandwich is heated to ∼75-80 °C to melt the parafilm, adhering the coverslips and creating the flow channel.

The following steps describe a click-chemistry procedure to covalently attach a DNA oligonucleotide to the slide. The reaction mixture consists of a 10 % mPEG-SVA (Laysan Bio: MPEG-SVA-5000) solution in 0.1 M potassium tetraborate pH 8.1. To that, DBCO-PEG4-NHS ester (Conju-Probe: CP-2028) at 400 μM resuspended in anhydrous DMSO (ThermoFisher Scientific: D12345) is combined at a 40:1 ratio with a 10 μM azide modified oligo (Sigma-Aldrich) for a final concentration of 50 nM DNA in the reaction mixture. The reaction mixture is injected into the flow cell and incubated for 1 hour at 23 °C. It is subsequently rinsed from the flowcell with Milli-Q water and the same reaction and incubation is performed a second time using fresh reagents. After the second incubation, the flow cell is rinsed thoroughly with Milli-Q water and allowed to dry over 18-24 hours at 23 °C in a nitrogen storage box. After the flowcells are dry, they are individually vacuum sealed and frozen at −20 °C until the day of the experiment.

On the day of the experiment, one flowcell is removed and brought to room temperature before removal from the packaging. The surface is passivated with Blocking Reagent (Roche: 11096176001) resuspended in 0.5 xTE and allowed to incubate for 20 minutes before rinsing with the experiment buffer (0.5 xTE with 100 mM NaCl, 10 mM MgCl_2_, and 0.5 % Tween20). Separately, the ND is annealed to the tether by incubating at 45 °C for 1 hour and then cooled at 1 °C/min. The annealing is checked using AGE (**Supplementary Fig. S15**). The ND+tether construct is then incubated with streptavidin coated superparamagnetic particles (prerinsed in experiment buffer) (Thermo-Scientific: Dynabeads M-280) for 15 minutes at room temperature. Pre-rinsed 4.5 µm streptavidin labeled polystyrene particles (Spherotech SVP-40-5) are combined with 1 µM biotinylated oligoes that are a reverse complement to the oligo covalently attached to the slide and incubated for 20 minutes at room temperature. Excess DNA is washed from the sample by repeated centrifugal pelleting, supernatant removal, and resuspension. These beads will adhere to the slide surface through many DNA’s on the bead surface base pairing with the DNA on the slide surface and act as a fiduciary mark during the experiment. Just before injection onto the slide, the ND+tether+Dynabead construct is combined with the DNA coated fiduciary particles.

### Single molecule force spectroscopy experiments on a magnetic tweezer system

Experiments were performed on a home-built magnetic tweezer system using an Olympus IX-70 inverted microscope body. Data collection for multiple samples is collected in parallel with an in-house written LabVIEW program. Prior to data collection, each bead position is individually calibrated so that relative position change of each bead can be measured. The force-extension is performed by moving a pair of permanent neodymium magnets close to the sample at a rate of 0.1 mm/s. The magnets are then retracted at the same rate. This process can be repeated many times to collect multiple opening events from the same tethered sample.

The force exerted on each molecule is calculated using the equipartition theorem and the approximation of the bead-tether system as an inverted pendulum. These two concepts result in the equation:

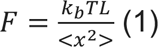

Where *k_b_* is the Boltzmann constant, *T* is the temperature, *L* is end-to-end distance of the molecule, and *x* is the deviation of the bead from the central position due to Brownian fluctuations.

### Modeling the NanoDyn as a torsional spring

To fit the experimental force extension curves, a model of the mechanical behavior of the ND+tether system was created. The model uses a worm-like chain for the tether, attached on one end to the surface and at the other end to a torsional spring, which represents the ND and its attachment to the bead. To avoid numerical artifacts due to the finite maximal extension of this model, we added a soft constraint on the physical length of the ND. We can then fit the force-extension curve predicted by the model to the experimental force-extension curves, by varying the four parameters of the model, namely the unknown offset in the distance measurements between the surface and the bead, the length extension constant of the ND, the spring constant of the torsional spring, and the initial angle Ɵ_o_, of the torsional spring with respect to the surface attachment point of the tether. A diagram of the system can be found in **Supplementary Fig. S7**. To account for the rupture, we first subtract the rupture from the high force regime data to give us a continuous data set to fit our low regime model to. We can then use the force point when the rupture occurred and the change in extension relative to the total length of the tether to algebraically calculate the newly acquired additional length of the system, adding that to the fitting process to give us our high force regime fit. A table of the fit parameters can be found in **Supplementary Fig. S7**. A more detailed summary of the fitting algorithm can be found in the **Supplementary Methods**.

### Flow incorporation of zippers into the ND in real-time

The setup of this experiment starts as previously described in the above methods. In the case of real-time zipper addition to the ND, a sample is prepared with the blocking strands omitted from the desired loops. To facilitate flow, the flowcell is attached via peristaltic pump tubing to a 1 mL syringe in a syringe pump (SyringePump.com). The pump is run at a rate of 10 µL/min to draw sample through the ∼30 µL flowcell sample chamber which is fed by a reservoir. To incorporate a zipper, 200 µL of 800 nM zipper oligonucleotide is added to the reservoir and drawn through the sample chamber for 7 minutes with the magnets exerting a force of ∼5 pN on the sample tethers, followed by a 20 minute incubation at ∼1 pN. During incubation, the sample reservoir is washed of remaining zipper by buffer exchanging 5 times with experiment buffer. Following the incubation, excess zipper is removed from the flowcell by drawing experiment buffer from the reservoir for 7 minutes with ∼5 pN force applied to the sample.

### Sample imaging by transmission electron microscopy (TEM)

For TEM imaging, structures were stained using a 1% Uranyl acetate solution and imaged on a FEI Tecnai G2 Spirit electron microscope. For each preparation, 6-8 μl of 1 nM DNA origami sample was wicked onto a glow-discharge-cleaned copper grid (Electron Microscopy Sciences, Hatfield, PA) and incubated for 5-10 minutes. The sample solution was then removed carefully with Whatman #4 filter paper and the grids were immediately stained with two 6 μl drops of 1 % Uranyl acetate solution. Grids were dried for at least 10 minutes before imaging.

### Log-likelihood removal of data from ND devices indicated as having incomplete zipper incorporation

We begin by using the 1-zipper distributions as reference models to determine if data from a single ND designed to have 2 zippers is “better fit” by the 1-zipper model. The 1-zipper distribution is assumed to be “correct” given the presence of the opening events. For each 2-zipper ND, the following calculation is performed:

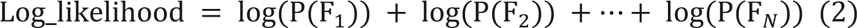

Where *P*(*F*) is the normalized probability of an opening force, *F*, being in a given distribution. *F* ranges from 1 to N where N is the total number of opening events for that device. To account for 0 count bins in the probability distributions, we added +1 pseudo-counts to each bin and renormalized the distributions accordingly. Not doing so results in undefined terms in the summation. This calculation is performed three times; once for each of the 1-zipper distributions (which are the 2 individual zippers contained in the 2-zipper ND) and once for the 2-zipper distribution. Smaller probabilities are more heavily weighted towards larger negative values, so whichever of the 3 calculated values results in the least negative number (closer to zero) is declared to be the distribution best fitting that list of forces. After testing all 2-zipper devices in this way, any of the 2-zipper devices indicated as being better fit by a 1-zipper distribution was removed from the aggregate data. We then repeated the same process for the 3-zipper ND data using the three 1-zipper distributions, the ‘cleaned-up’ 2-zipper distribution, and the 3-zipper distributions, for a total of 5 outputs. Again, any of the 3-zipper devices indicated as only having 1 or 2 zippers, were removed from the aggregate data. Histograms showing the data removal can be found in **Supplementary Fig. S8**.

### Estimate of the cumulative probability distribution for a ND with multiple force-sensing loops

To estimate the expected cumulative probability distribution (CPD) for a ND containing multiple force-sensing loops we added the CPD’s of the individual force-sensing loops. We assume that the force, *F*, is divided equally across all force-sensing loops and that once one of the loops is opened, they all open due to the increase in force on the remaining zipper(s). This is supported by experimental data in that we do not see multiple smaller opening steps during force loading. For a ND with 2 force-sensing loops, we calculate the cumulative probability that both zippers release at *F*, which is equivalent to the cumulative probability, *P_c_*, of one zipper or the other breaking at *F*/2 given our assumptions. This results in the equation:

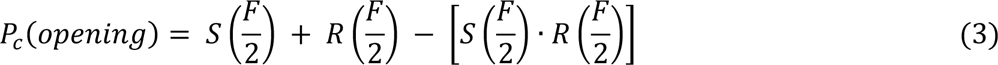

Where *S* and *R* are the CPD’s for the two individual force-sensing loops. We use a similar line of thinking for the ND with three force-sensing loops where *F* is distributed evenly across three zippers. This results in:

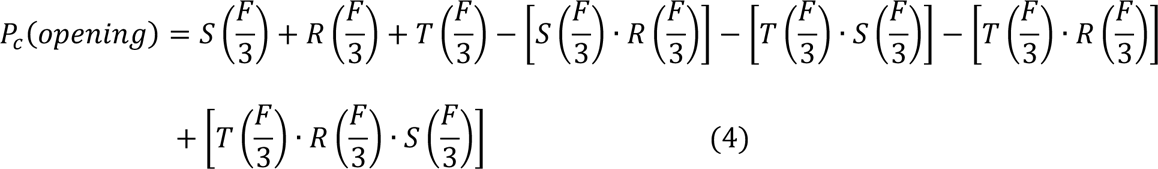

*T* is the CPD for a third force-sensing loop.

## Data Availability statement

The experimental data sets are either included in this submission, the supplemental information, or are available from the authors upon request.

## ACKNOWLEDGEMENTS

We are grateful to the Poirier, Castro, Bundschuh and Winter Lab members for insightful discussions and feedback on this work. We thank Yi Luo for help with developing our slide functionalization protocol. Transmission electron microscopy images were acquired at the OSU Campus Microscopy and Imaging Facility, which is supported in part by grant number P30 CA016058, National Cancer Institute, Bethesda, MD. Department of Energy [DESC0017270 to M.G.P, C.E.C, J.O.W.], National Science Foundation [CCMI1921881 to M.G.P., C.E.C.; DMR1719316 to R.A.B.]; National Institutes of Health [R35 GM139564 to M.G.P.]; Funding for open access charge: Department of Energy.

## AUTHOR CONTRIBUTIONS

A.R. prepared the DO samples, DNA tethers, and flow cells, carried out force spectroscopy measurements, and data analysis; H.H. developed the TorWLC model; M.N. carried out the TEM imaging; PB provided support for the DO sample preparation; M.G.P, R.B., and C.E.C. supervised the study and helped with data interpretation; A.R. and M.G.P. wrote the manuscript; All authors edited the manuscript and approved the final version.

## REFERENCES

1. Wang, Y. et al. Force-Dependent Interactions between Talin and Full-Length Vinculin. J. Am. Chem. Soc. 143, 14726–14737 (2021).

2. Assad, J. A., Shepherd, G. M. G. & Corey, D. P. Tip-link integrity and mechanical transduction in vertebrate hair cells. Neuron 7, 985–994 (1991).

3. Schnitzer, M. J., Visscher, K. & Block, S. M. Force production by single kinesin motors. Nat. Cell Biol. 2, (2000).

4. Ke, C., Humeniuk, M., S-Gracz, H. & Marszalek, P. E. Direct measurements of base stacking interactions in DNA by single-molecule atomic-force spectroscopy. Phys. Rev. Lett. 99, 1–4 (2007).

5. Lavelle, C., Victor, J. & Zlatanova, J. Chromatin Fiber Dynamics under Tension and Torsion. 1557–1579 (2010) doi:10.3390/ijms11041557.

6. Neuman, K. C. & Nagy, A. Single-molecule force spectroscopy : optical tweezers, magnetic tweezers and atomic force microscopy. 5, 491–505 (2008).

7. Abbondanzieri, E. A., Greenleaf, W. J., Shaevitz, J. W., Landick, R. & Block, S. M. Direct observation of base-pair stepping by RNA polymerase. Nature 438, 460–465 (2005).

8. Neuman, K. C. & Block, S. M. Optical trapping. Review of Scientific Instruments vol. 75 2787–2809 (2004).

9. Bausch, A. R., Möller, W. & Sackmann, E. Measurement of Local Viscoelasticity and Forces in Living Cells by Magnetic Tweezers. Biophys. J. 76, 573–579 (1999).

10. Hinterdorfer, P. & Dufrêne, Y. F. Detection and localization of single molecular recognition events using atomic force microscopy. Nat. Methods 3, 347–355 (2006).

11. Wojcikiewicz, E. P., Zhang, X. & Moy, V. T. Force and compliance measurements on living cells using atomic force microscopy (AFM). Biol. Proced. Online 6, 1–9 (2004).

12. Roy Chowdhury, S., Cao, J., He, Y. & Peter Lu, H. Revealing Abrupt and Spontaneous Ruptures of Protein Native Structure under picoNewton Compressive Force Manipulation. ACS Nano 12, 2448–2454 (2018).

13. Lee, G. U., Kidwell, D. A. & Colton, R. J. Sensing Discrete Streptavidin–Biotin Interactions with Atomic Force Microscopy. Langmuir 10, 354–357 (1994).

14. Tateishi-Karimta, H. & Sugimoto, N. Control of stability and structure of nucleic acids using cosolutes. Methods 67, 151–158 (2014).

15. Pielak, G. J. & Miklos, A. C. Crowding and function reunite. Proc. Natl. Acad. Sci. U. S. A. 107, 17457–17458 (2010).

16. Yamamoto, K., Ota, N. & Tanaka, Y. Nanofluidic Devices and Applications for Biological Analyses. Anal. Chem. 93, 332–349 (2021).

17. Abgrall, P. & Trung Nguyenk, N. Nanofluidic Devices and Their Applications. Anal. Chem. 80, 155–214 (2008).

18. Zhang, Q. et al. DNA origami as an in vivo drug delivery vehicle for cancer therapy. ACS Nano 8, 6633–6643 (2014).

19. Jiang, Q., Liu, S., Liu, J., Wang, Z. G. & Ding, B. Rationally Designed DNA-Origami Nanomaterials for Drug Delivery In Vivo. Adv. Mater. 31, 1–6 (2019).

20. Ge, Z. et al. DNA Origami-Enabled Engineering of Ligand–Drug Conjugates for Targeted Drug Delivery. Small 16, 6–11 (2020).

21. Wang, S. et al. DNA Origami-Enabled Biosensors. Sensors 1–18 (2020) doi:doi:10.3390/s20236899.

22. Pal, S., Naik, A., Rao, A., Chakraborty, B. & Varma, M. M. Aptamer-DNA Origami-Functionalized Solid-State Nanopores for Single-Molecule Sensing of G-Quadruplex Formation. ACS Appl. Nano Mater. 5, 8804–8810 (2022).

23. Nickels, P. C. et al. Molecular force spectroscopy with a DNA origami-based nanoscopic force clamp. Science (80-.). 354, 305–307 (2016).

24. Kramm, K. et al. DNA origami-based single-molecule force spectroscopy elucidates RNA Polymerase III pre-initiation complex stability. Nat. Commun. 11, 1–12 (2020).

25. Funke, J. J. et al. Uncovering the forces between nucleosomes using DNA origami. Sci. Adv. 2, (2016).

26. Le, J. V. et al. Probing Nucleosome Stability with a DNA Origami Nanocaliper. ACS Nano 10, 7073–7084 (2016).

27. Kilchherr, F. et al. Single-molecule dissection of stacking forces in DNA. Science (80-.). 353, (2016).

28. Pfitzner, E., et al. Rigid DNA beams for high-resolution single-molecule mechanics. Angew. Chemie - Int. Ed. 52, 7766–7771 (2013).

29. Dutta, P. K. et al. Programmable Multivalent DNA-Origami Tension Probes for Reporting Cellular Traction Forces. Nano Lett. 18, 4803–4811 (2018).

30. Hudoba, M. W., Luo, Y., Zacharias, A., Poirier, M. G. & Castro, C. E. Dynamic DNA Origami Device for Measuring Compressive Depletion Forces. ACS Nano 11, 6566–6573 (2017).

31. Darcy, M. et al. High-Force Application by a Nanoscale DNA Force Spectrometer SI. ACS Nano 16, 5682–5695 (2022).

32. Wang, Y. et al. A nanoscale DNA force spectrometer capable of applying tension and compression on biomolecules. Nucleic Acids Res. 49, 8987–8999 (2021).

33. Marras, A. E., Zhou, L., Su, H.-J. & Castro, C. E. Programmable motion of DNA origami mechanisms. Proc. Natl. Acad. Sci. 112 VN-, 713–718 (2015).

34. Zhou, L., Marras, A. E., Su, H. J. & Castro, C. E. Direct design of an energy landscape with bistable DNA origami mechanisms. Nano Lett. 15, 1815–1821 (2015).

35. Ijäs, H., Hakaste, I., Shen, B., Kostiainen, M. A. & Linko, V. Reconfigurable DNA Origami Nanocapsule for pH-Controlled Encapsulation and Display of Cargo. ACS Nano 13, 5959–5967 (2019).

36. Castro, C. E. et al. A primer to scaffolded DNA origami. Nat. Methods 8, 221–229 (2011).

37. Shih, W. M., Quispe, J. D. & Joyce, G. F. A 1.7-kilobase single-stranded DNA that folds into a nanoscale octahedron. Nature 427, 618–621 (2004).

38. Rothemund, P. W. K. Folding DNA to create nanoscale shapes and patterns. Nature 440, 297–302 (2006).

39. Woodside, M. T. et al. Nanomechanical measurements of the sequence-dependent folding landscapes of single nucleic acid hairpins. Proc. Natl. Acad. Sci. U. S. A. 103, 6190–6195 (2006).

40. Mossa, A., Manosas, M., Forns, N., Huguet, J. M. & Ritort, F. Dynamic force spectroscopy of DNA hairpins: I. Force kinetics and free energy landscapes. J. Stat. Mech. Theory Exp. 2009, (2009).

41. Lee, C. H., Danilowicz, C., Conroy, R. S., Coljee, V. W. & Prentiss, M. Impacts of magnesium ions on the unzipping of λ-phage DNA. J. Phys. Condens. Matter 18, (2006).

42. Nakano, S. I., Fujimoto, M., Hara, H. & Sugimoto, N. Nucleic acid duplex stability: Influence of base composition on cation effects. Nucleic Acids Res. 27, 2957–2965 (1999).

43. Bockelmann, U., Thomen, P., Essevaz-Roulet, B., Viasnoff, V. & Heslot, F. Unzipping DNA with optical tweezers: High sequence sensitivity and force flips. Biophys. J. 82, 1537–1553 (2002).

44. Woo, S. & Rothemund, P. W. K. Programmable molecular recognition based on the geometry of DNA nanostructures. Nat. Chem. 3, 620–627 (2011).

45. Huang, C. M., Kucinic, A., Le, J. V., Castro, C. E. & Su, H. J. Uncertainty quantification of a DNA origami mechanism using a coarse-grained model and kinematic variance analysis. Nanoscale 11, 1647–1660 (2019).

46. Ji, J., Karna, D. & Mao, H. DNA origami nano-mechanics. Chem. Soc. Rev. 50, 11966–11978 (2021).

47. Zhou, L., Marras, A. E., Su, H. J. & Castro, C. E. DNA origami compliant nanostructures with tunable mechanical properties. ACS Nano 8, 27–34 (2014).

48. Rickgauer, J. P. et al. Portal Motor Velocity and Internal Force Resisting Viral DNA Packaging in Bacteriophage f29. Biophys. J. 94, 159–167 (2008).

49. Chemla, Y. R. et al. Mechanism of Force Generation of a Viral DNA Packaging Motor. 122, 683–692 (2006).

50. Poirier, M. G. & Marko, J. F. Mitotic chromosomes are chromatin networks without a mechanically contiguous protein scaffold. 99, 15393–15397 (2002).

51. Michel Grandbois, Martin Beyer, Matthias Rief, Hauke Clausen-Schaumann, H. E. G. How Strong Is a Covalent Bond? Science (80-.). 283, 1727–1730 (1999).

52. Roein-peikar, M., Xu, Q., Wang, X. & Ha, T. Ultrasensitivity of Cell Adhesion to the Presence of Mechanically Strong Ligands. 011001, 1–9 (2016).

53. Jo, M. H., Cottle, W. T. & Ha, T. Real-Time Measurement of Molecular Tension during Cell Adhesion and Migration Using Multiplexed Differential Analysis of Tension Gauge Tethers. ACS Biomater. Sci. Eng. 5, 3856–3863 (2019).

54. Stahl, E., Martin, T. G., Praetorius, F. & Dietz, H. Facile and scalable preparation of pure and dense DNA origami solutions. Angew. Chemie - Int. Ed. 53, 12735–12740 (2014).

55. Xin, Y. et al. Cryopreservation of DNA Origami Nanostructures. Small 16, (2020).

56. Luo, Y. et al. Resolving Molecular Heterogeneity with Single-Molecule Centrifugation. J. Am. Chem. Soc. (2022) doi:10.1021/jacs.2c11450.

57. Chandradoss, S. D. et al. Surface passivation for single-molecule protein studies. J. Vis. Exp. 1–8 (2014) doi:10.3791/50549.

