## Supplementary Information for "Cooperative control of a DNA origami force sensor"

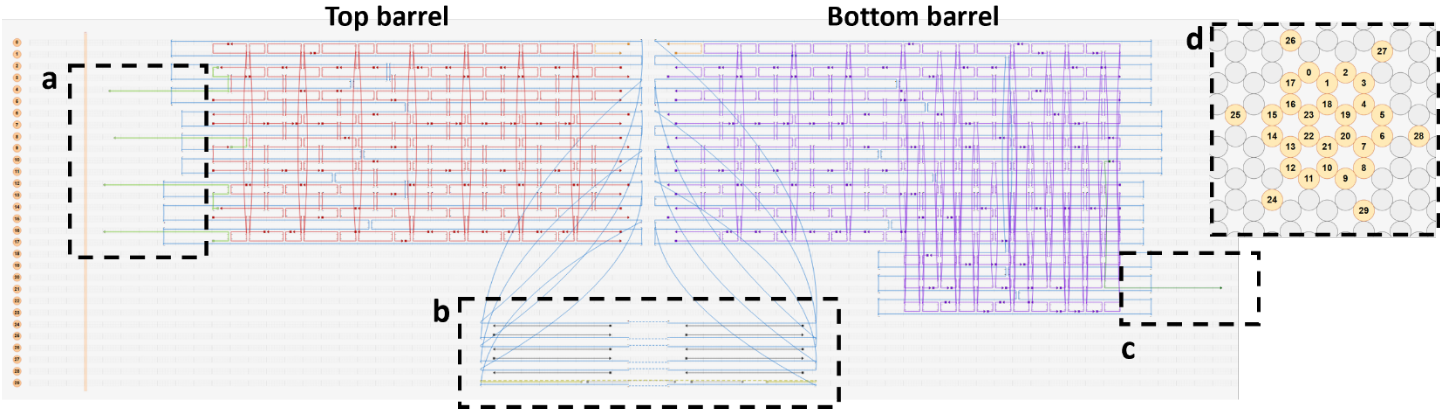


**Supplementary Figure S1.** caDNAno drawing of the DNA origami NanoDyn (ND): (**a**) 4, 24 nt ssDNA overhangs protruding from helixes 4, 8, 12, and 16 which anneal to a 25 nt ssDNA biotinylated oligonucleotide. There is a 22 bp overlap, leaving a total of 5 nt's unpaired spacers. (**b**) 6, 116 nt sections of ssDNA scaffold sequences attaching the top and bottom barrels of the ND referred to in the main text as 'loops". (**c**) 30 nt ssDNA overhang protruding from helix 21 of the ND for attachment to the DNA tether handle. (**d**) Cross-sectional view of the bottom barrel of the ND. The 6 helixes 24-29 indicate the routing of the loops outside the main body of the structure.

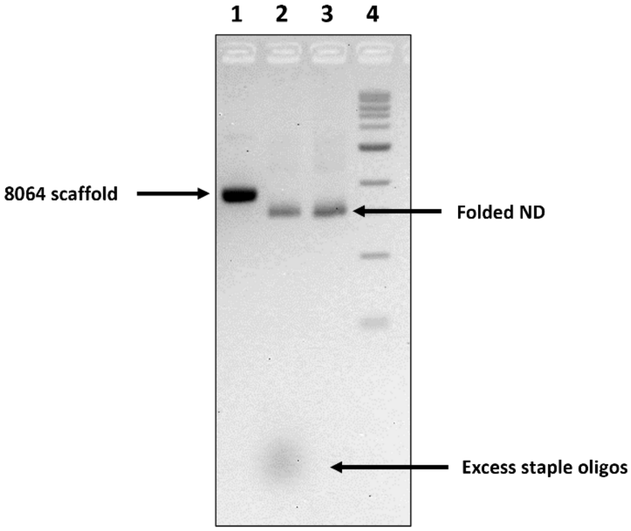


**Supplementary Figure S2.** Example of a NanoDyn fold and purification of the base version of the device (no zippers). **(1)** 8064 nt ssDNA scaffold **(2)** Unpurified ND omitting the L6 and L3 zippers. **(3)** PEG purified ND omitting the L6 and L3 zippers. **(4)** 1 kb DNA ladder.

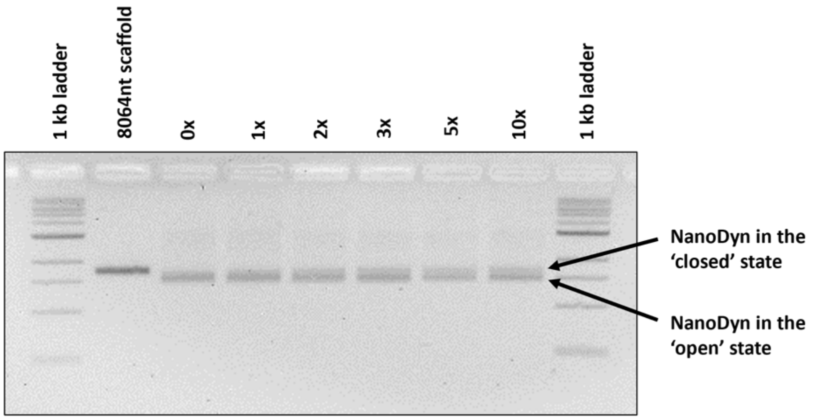


**Supplementary Figure S3.** Representative ND zipper incorporation gel for L6-15nt. From left to right, the L6-15 nt zipper was titrated from 0x to 10x to determine the optimal zipper concentration. The lower band is ND that either has no zipper incorporated, or has a zipper, but in the course of electrophoresis has not remained closed. The upper band is ND that has the zipper incorporated and remained closed.

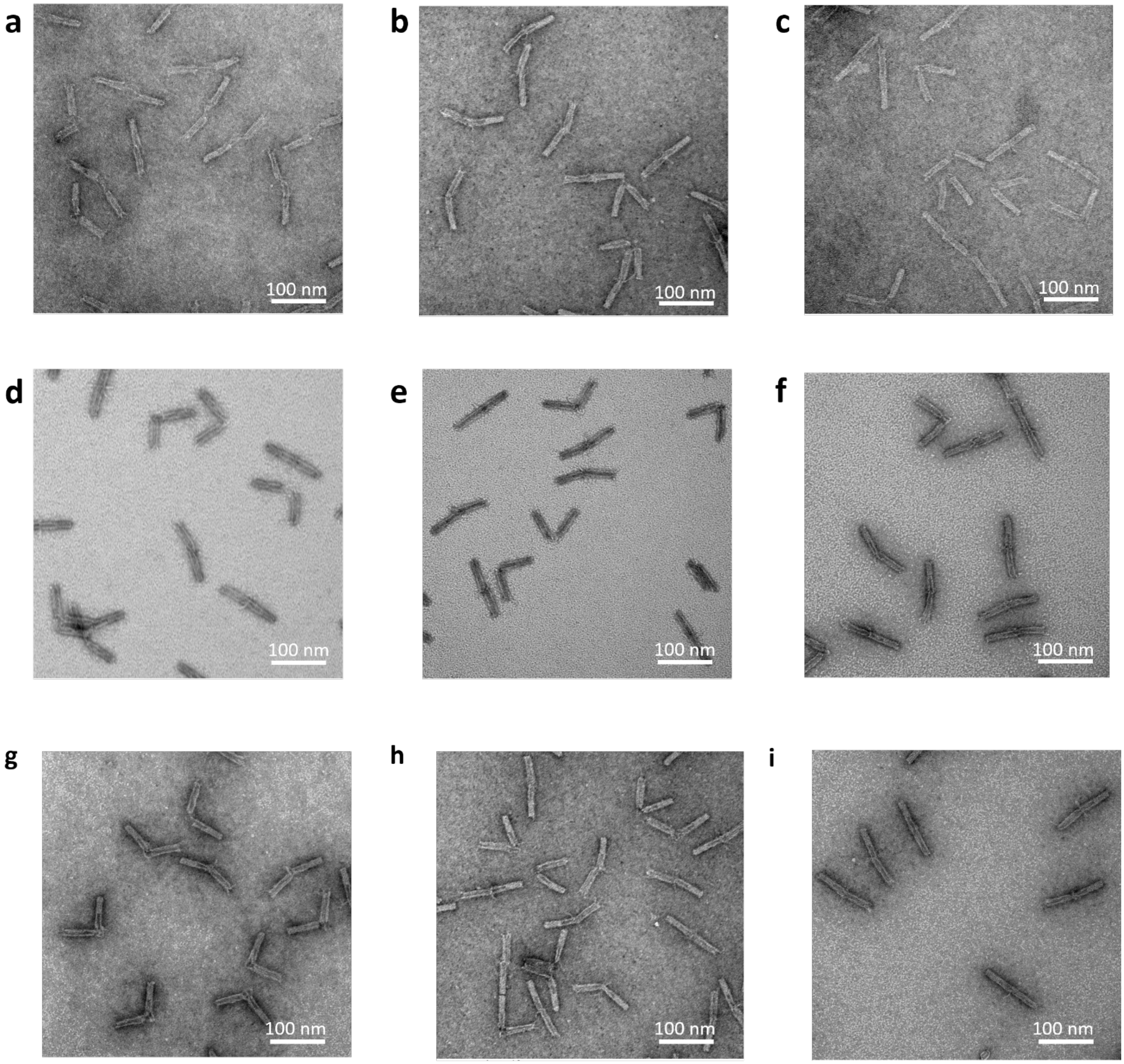


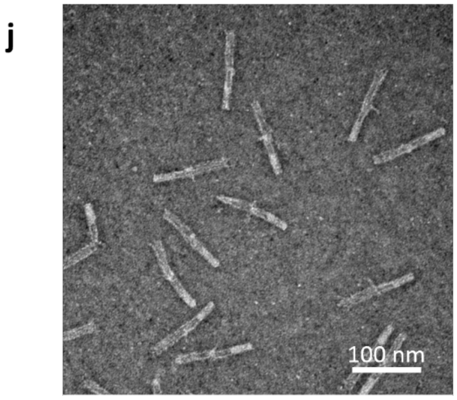


**Supplementary Figure S4.** Representative TEM imagine of ND's with various zippers incorporated. (**a**) No zippers, completely unconstrained. (**b**) L6-13nt. (**c**) L6-15nt. (**d**) L6-17nt. (**e**) L6-19nt (**f**) L6-21nt (**g**) L3-11nt. (**h**) L1-13nt. (**i**) L6+L3. (**j**) L6+L3+L1.

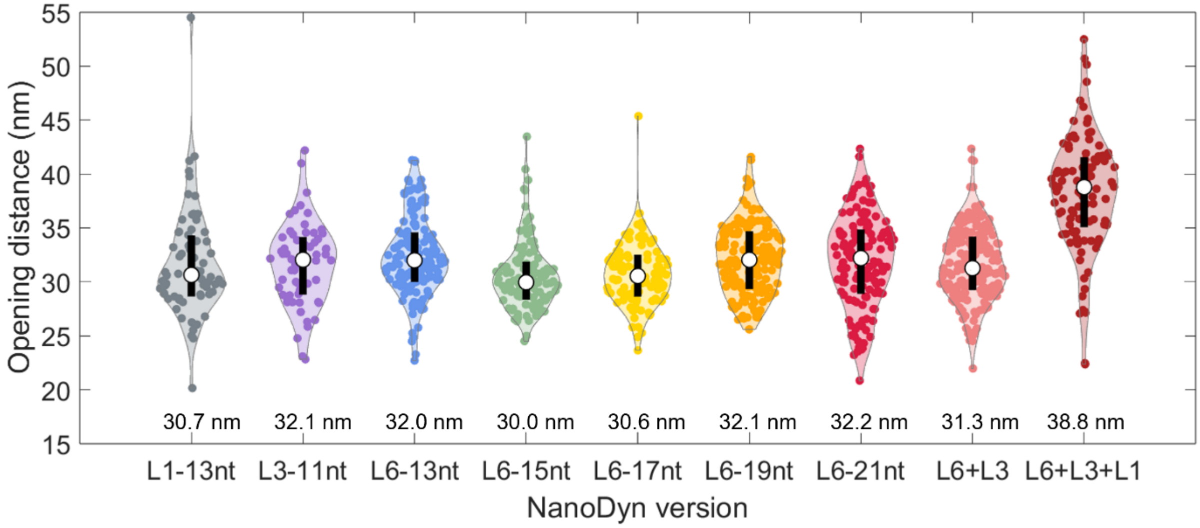


**Supplementary Figure S5.** The opening distance distribution for each ND version: Each colored dot represents the opening distance for each opening event. The white dot indicates the median opening distance which is listed below for each device version and the black bar is the first quartile above and below the median. The median opening distance, mean, SD, SEM, and interquartile values can be found in **Supplementary Table S2**. The L6-13nt + L3-11nt (pink) and L6-13nt + L3-11nt + L1-13nt (dark red) device names have been abbreviated L6+L3 and L6+L3+L1 respectively. The L6+L3+L1 distribution had 4 low distance outliers removed as the opening event occurred at the end of the data collection period and thus, an accurate opening distance could not be determined.

**a**

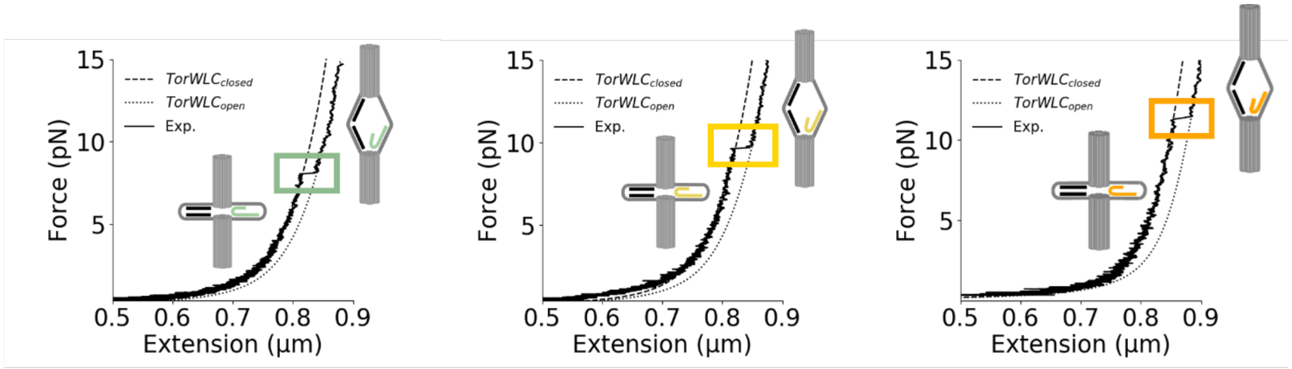


**b**

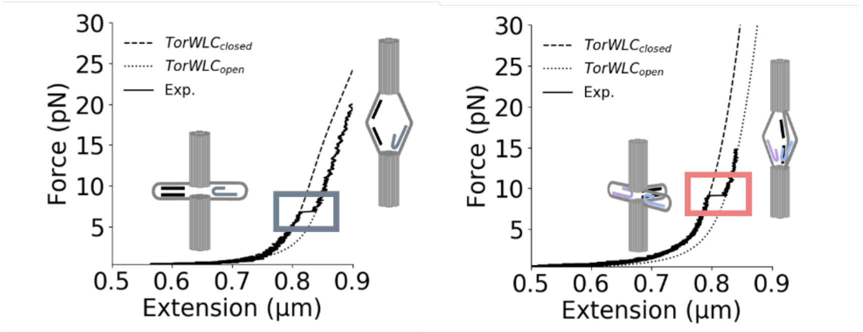


**Supplementary Figure S6:** Zippers of variable length and number modulate the opening force: Force extension data was fit to a Torsional spring + worm-like chain model (TorWLC) for both the closed ($TorWLC_{closed}$) and open ($TorWLC_{open}$) states. (**a**) Representative force extension curves for L6-15nt (green), L6-17nt (yellow), L6-19nt (orange). (**b**) Representative force extension curves for L1-13nt (grey), L6+L3 (pink).

**a**

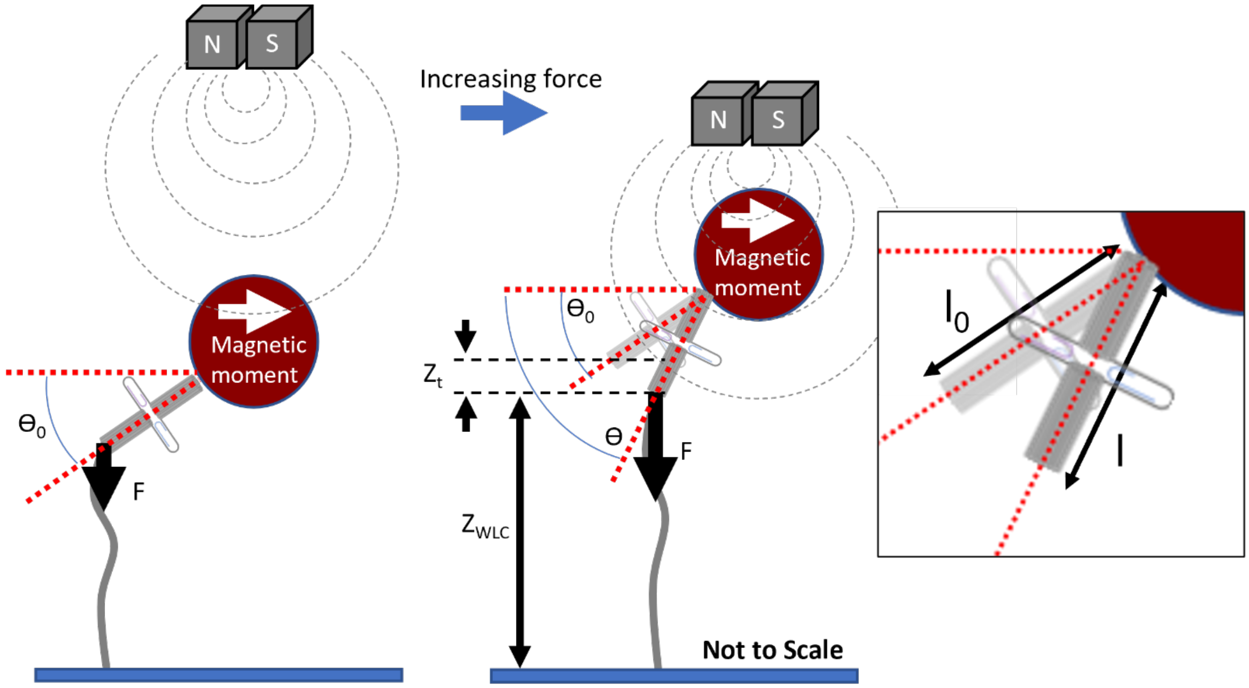


**b**

| Color ID | Device version | $K_l$ (pN/nm) | $K_\theta$ (pNnm/rad) | $\Theta_0$ (rad) |
|---|---|---|---|---|
| | L6-13nt | $8.89 \cdot 10^{3}$ | $1.12 \cdot 10^{4}$ | -0.06 |
| | L6-15nt | $1.05 \cdot 10^{4}$ | $1.90 \cdot 10^{3}$ | -0.16 |
| | L6-17nt | $9.42 \cdot 10^{3}$ | $2.07 \cdot 10^{3}$ | -0.18 |
| | L6-19nt | $6.72 \cdot 10^{3}$ | $3.73 \cdot 10^{3}$ | -0.40 |
| | L6-21nt | $1.18 \cdot 10^{3}$ | $1.97 \cdot 10^{3}$ | -1.11 |
| | L3-11nt | $8.98 \cdot 10^{3}$ | $2.85 \cdot 10^{3}$ | -0.02 |
| | L1-13nt | $7.25 \cdot 10^{3}$ | $2.56 \cdot 10^{3}$ | -1.18 |
| | L6+L3 | $5.76 \cdot 10^{3}$ | $6.02 \cdot 10^{3}$ | -0.04 |
| | L6+L3+L1 | $5.51 \cdot 10^{3}$ | $4.39 \cdot 10^{3}$ | -0.66 |
| | Flow incorp: no zipper | $1.30 \cdot 10^{4}$ | $5.17 \cdot 10^{2}$ | -0.61 |
| | Flow incorp: L6-13nt | $9.59 \cdot 10^{3}$ | $2.33 \cdot 10^{3}$ | -0.07 |
| | Flow incorp: L6+L3 | $8.02 \cdot 10^{3}$ | $2.99 \cdot 10^{3}$ | -0.02 |

**Supplementary Figure S7.** Torsional spring + Worm-Like Chain model (TorWLC): (**a**) This model was developed to account for variable torque applied to the ND based on the relative position to which the ND binds to the bead surface and the moment of the bead which will align with the applied magnetic field. The angle $\Theta$ will increase from its initial value $\Theta_o$ until the force vector, F, is parallel with the lever arm (the NanoDyn). $Z_{WLC}$ is the extension due to the Worm-Like Chain extension of the dsDNA tether and $Z_t$ is the extension due to the change in the angle $\Theta$ of the NanoDyn relative to the horizontal. $l_0$ is the initial length of the NanoDyn and l is the final length of the NanoDyn. (**b**) The fit parameters listed were obtained for each of the force extension plots in the main body of the paper. $K_l$ is a linear spring constant and $K_\theta$ is a torsional spring constant.

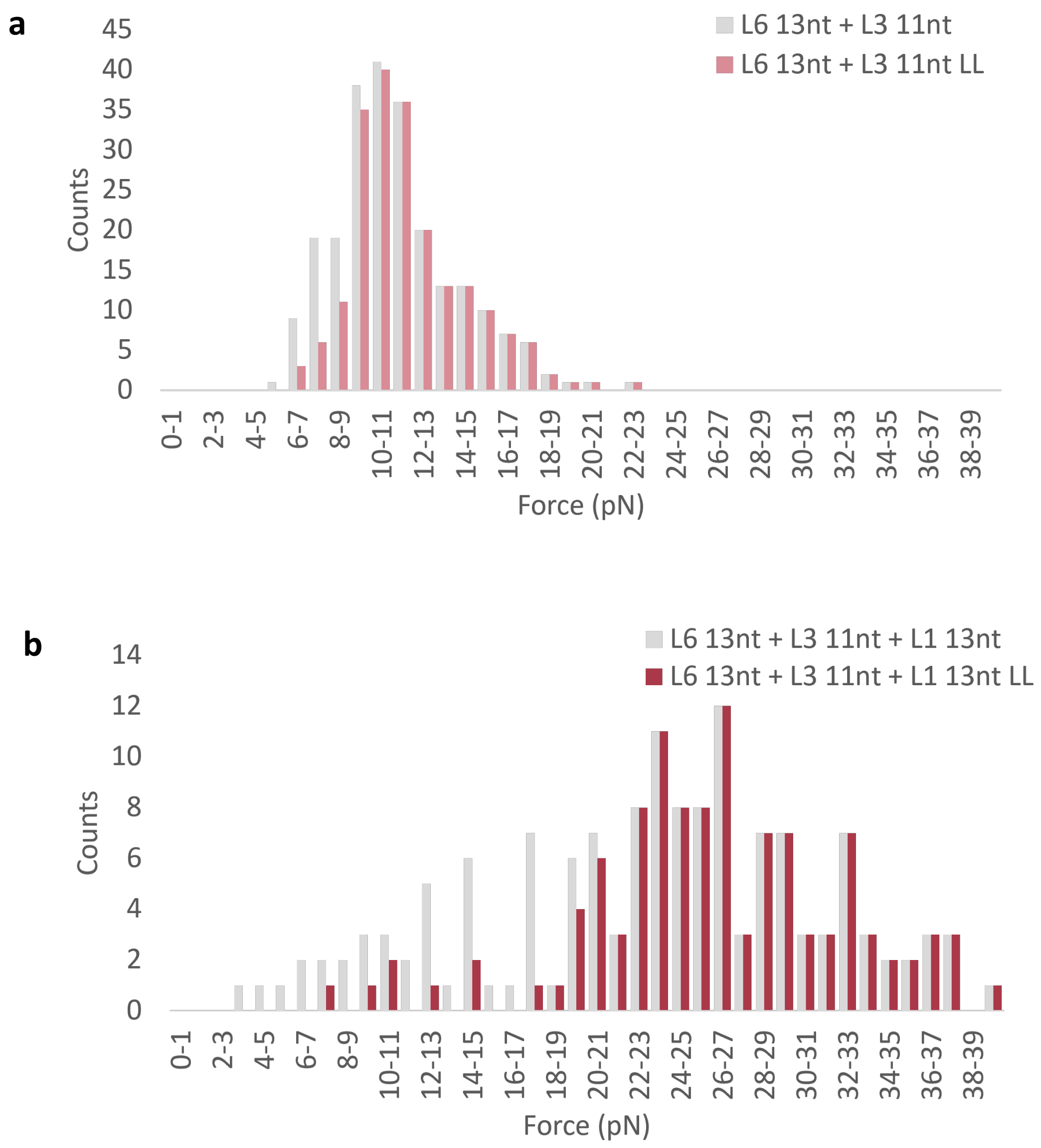


**Supplementary Figure S8.** Verification of zipper incorporation in the 2 and 3-zipper ND's using log-likelihood analysis. The data remaining after the log-likelihood analysis is denoted with "LL" in the legend. (**a**) Histogram of opening forces for the L6+L3 ND before (grey) and after (pink) log-likelihood cleanup. (**b**) Histogram of opening forces for the L6+L3+L1 ND before (grey) and after (dark red) log-likelihood cleanup.

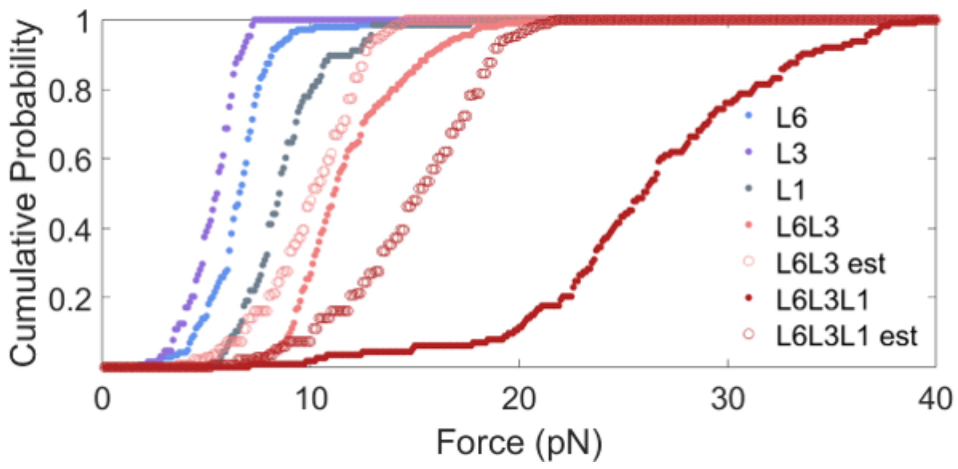


**Supplementary Figure S9.** Multi-zipper ND opening force estimation using cumulative sum addition. Experiment data are shown as solid dots and the estimates (est) as open circles. To estimate the L6+L3 opening force (pink) we summed the cumulative probability for the L6-13nt zipper (blue) and the L3-11nt zipper (purple). To estimate the L6+L3+L1 device (dark red) we summed the cumulative probability of L6-13nt zipper (blue), the L3-11nt zipper (purple), and the L1-13nt zipper (grey).

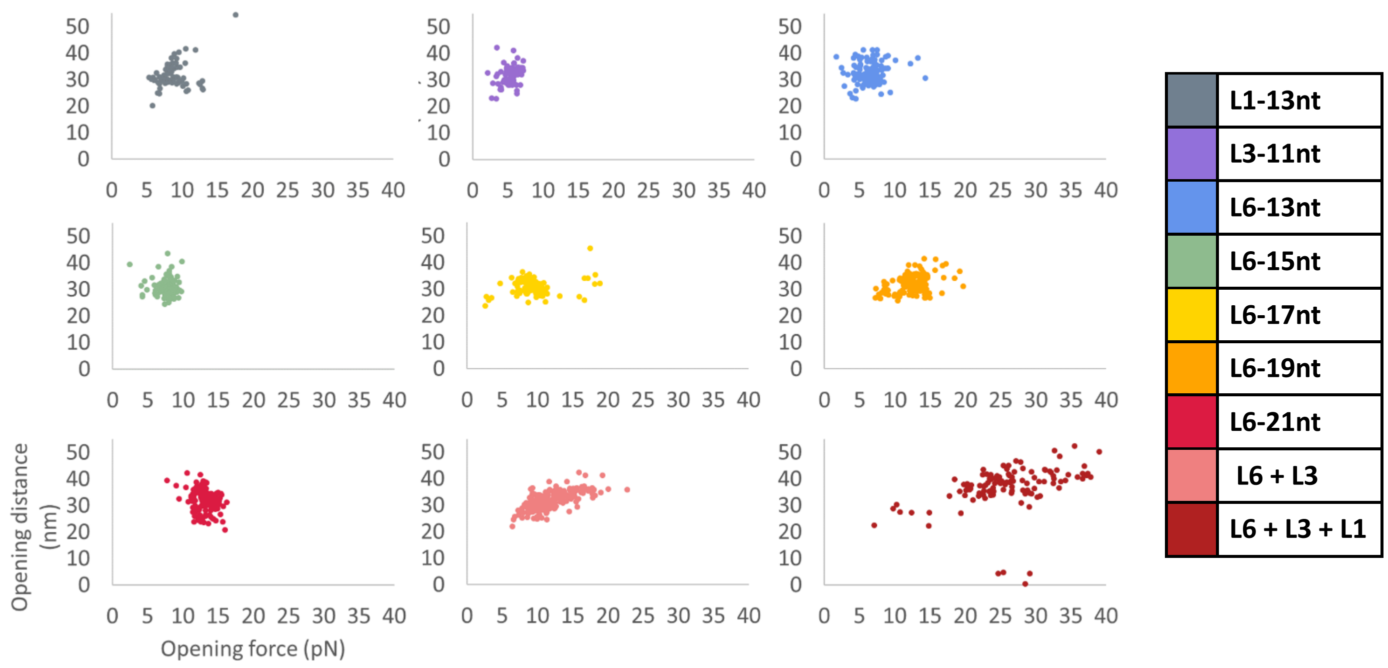


**Supplementary Figure S10:** For each opening event, the opening force vs opening distance was plotted for different ND versions.  Note that the 4 low-force point in the L6+L3+L1 plot were the outliers removed from the distribution in **Supplemental Fig S5**.

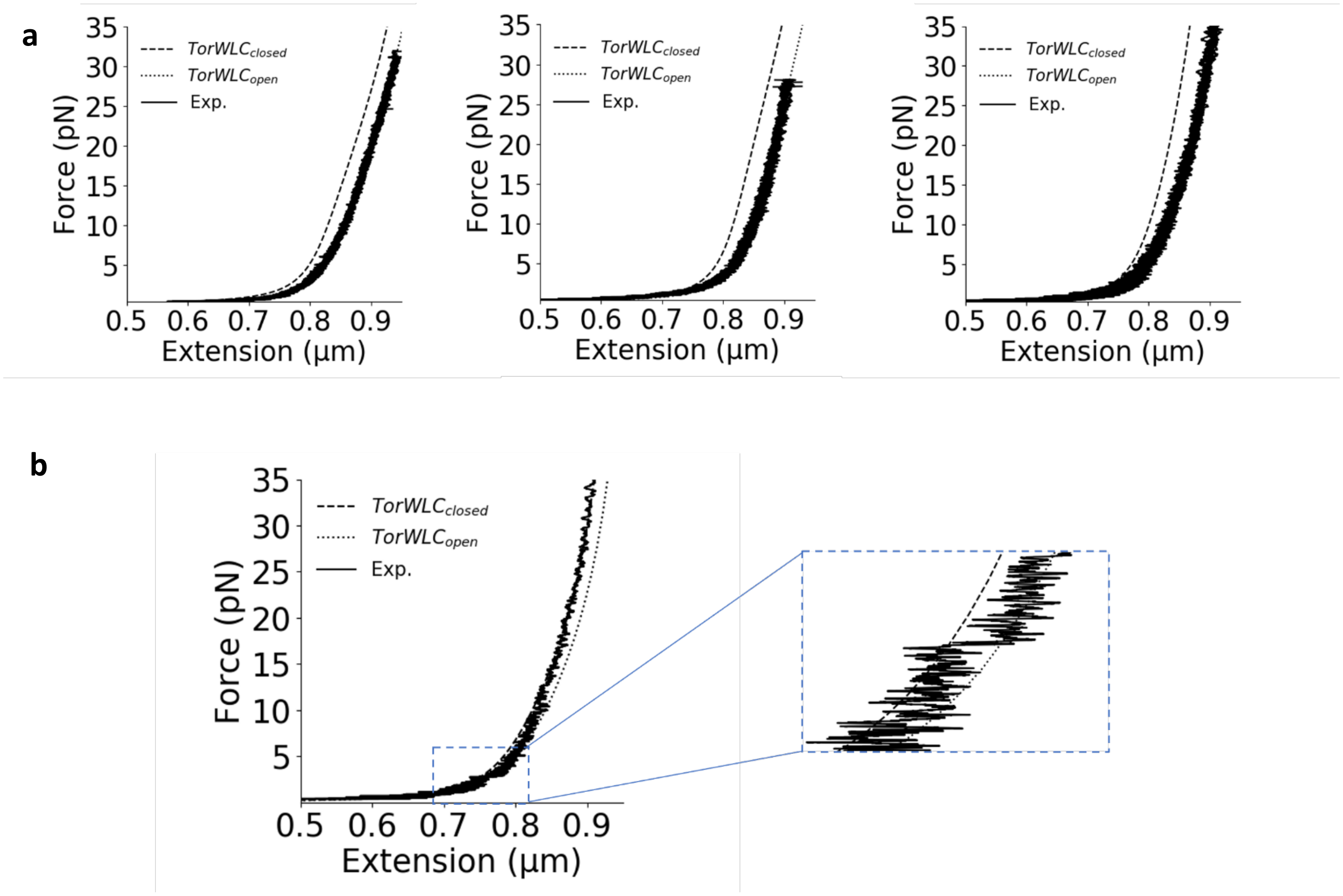


**Supplementary Figure S11.** Force extensions curves of the ND during unloading. (**a**) From left to right: 10 overlayed traces each of L1-13nt, L6-13nt, and L6+L3+L1, respectively. These unloading traces are the counterparts to the loading traces in main Figure 4. (**b**) Although relatively rare, closing events were occasionally observed. Most closures occurred at low force and are not easily distinguished in the bead fluctuation data. This event was collected from a L6+L3+L1, 3-zipper ND. Our fitting algorithm prioritizes fitting around the opening/closing step, thus, the closing event at low force skews the fit away from fitting the curve better at higher forces.

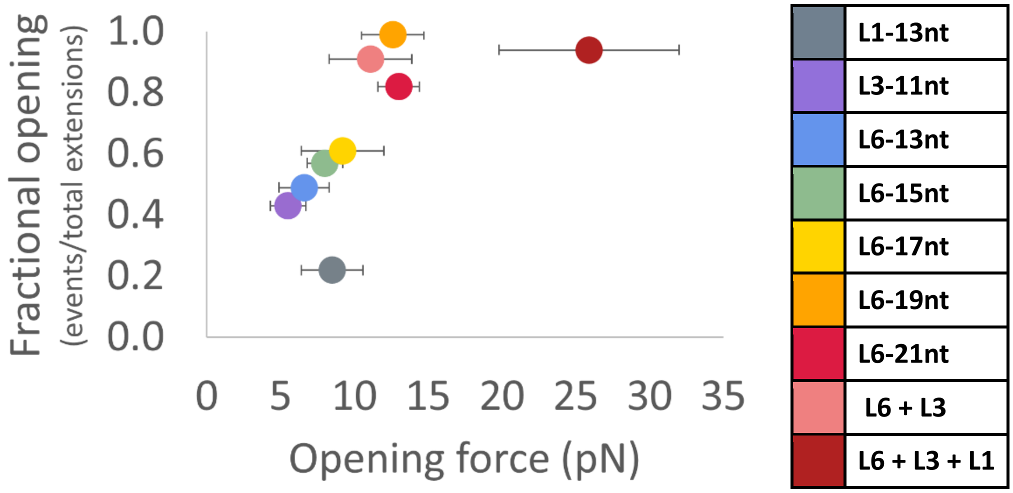


**Supplementary Figure S12:** Fractional opening of each ND version compared with its median opening force. The horizontal error bars are the standard deviation of the opening force. Numerical values can be found in **Supplementary Tables S1** and **S4.**

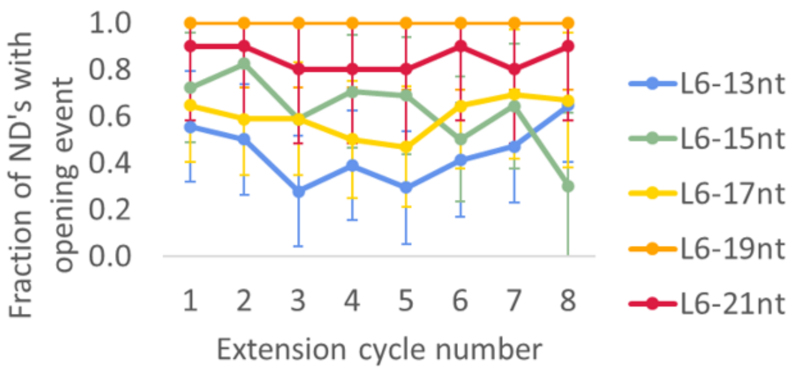


**Supplementary Figure S13:** Normalized fraction of ND openings per extension cycle. The standard error was calculated as 1/Sqrt(N) samples measured at cycle, M).

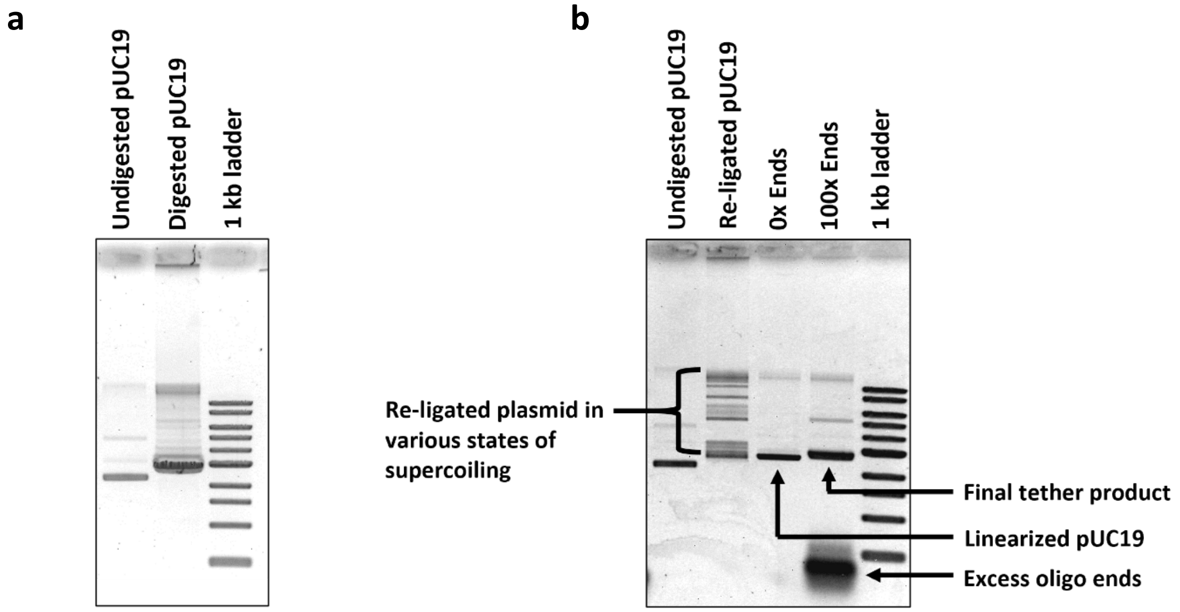


**Supplementary Figure S14.** Tether preparation for attaching the ND to the microscope slide. (**a**) Restriction enzyme BsaI was used to linearize a pUC19 plasmid. This leaves 4 nt sticky ends to which additional oligos are subsequently ligated. (**b**) 2 pairs of oligos are annealed to the linearized pUC19 sticky ends using T4 DNA ligase, resulting in a tether with a 30 nt overhang on one end and a 60 nt overhang on the other. We used 100x excess of the oligo ends for the final product, preventing recyclization and multimer formation of the pUC19 plasmid. This final product was then HPLC purified to remove excess ends (see **Methods**).

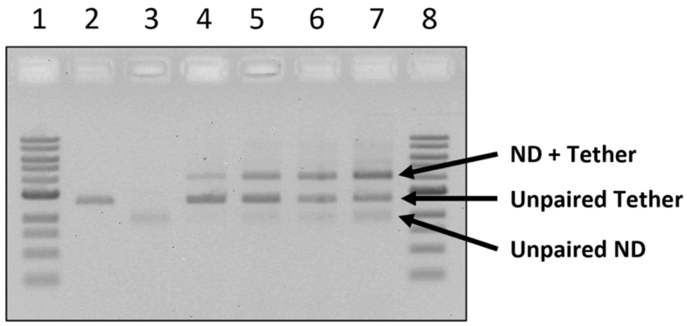


**Supplementary Figure S15.** Tether anneal to the ND: Tethers were annealed by titrating the amount of ND relative to the tether, which is held fixed. The ideal ratio maximizes attachment while minimizing free ND. From left to right: (**1**) 1 kb ladder. (**2**) dsDNA tether. (**3**) ND with L6-15nt zipper incorporated (**4**) 0.25x. (**5**) 0.5x. (**6**) 0.75x. (**7**) 1x. (**8**) 1 kb ladder.

| Color ID | NanoDyn version | Median opening force (pN) | Mean opening force ± SD (pN) | SEM (pN) | Interquartile (pN) | Total opening events measured | Total # of molecules studied |
|---|---|---|---|---|---|---|---|
| | **L1-13nt** | **8.5** | **8.7 ± 2.1** | **0.3** | **1.9** | **68** | **13** |
| | **L3-11nt** | **5.5** | **5.3 ± 1.2** | **0.1** | **1.6** | **64** | **11** |
| | **L6-13nt** | **6.6** | **6.5 ± 1.7** | **0.1** | **1.6** | **143** | **18** |
| | **L6-15nt** | **8.0** | **7.7 ± 1.2** | **0.1** | **1.1** | **113** | **11** |
| | **L6-17nt** | **9.2** | **9.5 ± 2.8** | **0.3** | **2.0** | **108** | **17** |
| | **L6-19nt** | **12.6** | **12.5 ± 2.1** | **0.2** | **2.5** | **162** | **13** |
| | **L6-21nt** | **13.0** | **13.1 ± 1.4** | **0.1** | **1.6** | **136** | **10** |
| | **L6 + L3** | **11.1** | **11.6 ± 2.8** | **0.2** | **3.3** | **205** | **23** |
| | **L6 + L3 + L1** | **25.9** | **26.0 ± 6.1** | **0.6** | **7.0** | **113** | **11** |

**Supplementary Table S1:** Table of measured forces, the associated error, and event counts for different ND versions.

| Color ID | NanoDyn version | Median opening distance (nm) | Mean opening distance ± SD (nm) | SEM (nm) | Interquartile: (nm) |
|---|---|---|---|---|---|
| | **L1-13nt** | **30.7** | **31.7 ± 4.9** | **0.6** | **5.0** |
| | **L3-11nt** | **32.1** | **31.7 ± 3.7** | **0.5** | **4.8** |
| | **L6-13nt** | **32.0** | **32.4 ± 3.6** | **0.3** | **4.1** |
| | **L6-15nt** | **30.0** | **30.6 ± 3.2** | **0.3** | **3.0** |
| | **L6-17nt** | **30.6** | **30.7 ± 2.9** | **0.3** | **3.4** |
| | **L6-19nt** | **32.1** | **32.2 ± 3.2** | **0.3** | **4.8** |
| | **L6-21nt** | **32.2** | **31.9 ± 4.2** | **0.4** | **5.4** |
| | **L6 + L3** | **31.3** | **31.6 ± 3.3** | **0.2** | **4.4** |
| | **L6 + L3 + L1** | **38.8** | **38.2 ± 5.2** | **0.5** | **6.0** |

**Supplementary Table S2:** Table of measured opening distances and the associated error for different ND versions.

| Color ID | Zipper name | Force-sensing strand anchor sequence + 5 nt poly-T spacer (5’-3’) | Zipper binding sequence (5’-3’) (*end connected to poly-T spacer) | Zipper melting temperature (°C) |
|---|---|---|---|---|
| | L6-13nt | TTTTTTCTGCGAACGAGTAGATTTAGTTTGACCAT | ATCAATTCTACTA* | 36.4 |
| | L6-15nt | TTTTTTCTGCGAACGAGTAGATTTAGTTTGACCAT | GCATCAATTCTACTA* | 46.0 |
| | L6-17nt | TTTTTTCTGCGAACGAGTAGATTTAGTTTGACCAT | TGGCATCAATTCTACTA* | 52.6 |
| | L6-19nt | TTTTTTCTGCGAACGAGTAGATTTAGTTTGACCAT | GGTGGCATCAATTCTACTA* | 57.2 |
| | L6-21nt | TTTTTTCTGCGAACGAGTAGATTTAGTTTGACCAT | AAGGTGGCATCAATTCTACTA* | 59.8 |
| | L3-11nt | CAAGAACGGGTATTAAACCAAGTACCGCTTTTT | *TAAGTCCTGAA | 35.1 |
| | L1-13nt | ATTAGACTTTACAAACAATTCGACAACTCGTATTTTTTT | *GGTTATCTAAAAT | 36.0 |

**Supplementary Table S3.** Table of the individual zipper designs used in the main text. The anchor sequence was designed such that the melting temperature was >60 °C. The 5nt poly-T was added so that there would not be a kink in the zipper oligo where the binding and anchor region meet. Melting temperatures were calculated using the IDT Oligo Analyzer Tool, using the experimental salt conditions of 100 mM NaCl and 10 mM $MgCl_2$. The oligo concentration was set to 40nM, which is the concentration used during post-fold incorporation of the zippers.

| Color ID | NanoDyn version | Total opening events / total extension cycles | | Median opening fraction | Mean opening fraction of individual NDs ± SD | SEM | Interquartile: |
|---|---|---|---|---|---|---|---|
| | L1-13nt | 68/304 | 22% | 0.13 | 0.28 ± 0.31 | 0.09 | 0.17 |
| | L3-11nt | 64/148 | 43% | 0.33 | 0.47 ± 0.29 | 0.09 | 0.31 |
| | L6-13nt | 143/292 | 49% | 0.43 | 0.46 ± 0.23 | 0.05 | 0.33 |
| | L6-15nt | 113/199 | 57% | 0.59 | 0.66 ± 0.27 | 0.06 | 0.52 |
| | L6-17nt | 108/178 | 61% | 0.60 | 0.56 ± 0.26 | 0.06 | 0.35 |
| | L6-19nt | 162/164 | 99% | 1.00 | 0.99 ± 0.04 | 0.01 | 0.00 |
| | L6-21nt | 136/166 | 82% | 1.00 | 0.83 ± 0.27 | 0.09 | 0.33 |
| | L6 + L3 | 205/225 | 91% | 1.00 | 0.92 ± 0.12 | 0.03 | 0.10 |
| | L6 + L3 + L1 | 113/120 | 94% | 1.00 | 0.95 ± 0.09 | 0.03 | 0.03 |

**Supplementary Table S4:** Table of opening fraction characteristics and the associated error for different ND versions.

**Supplementary Table S5.** Table of oligonucleotides used to fold the base NanoDyn structure.

| Name | Sequence |
|---|---|
| oligo1 | ATGCGTTCTAGCTGATAAACAGGAAATCCTGGAGGCCGCTAT |
| oligo2 | TAATGAGAGATCTACAATCAATCAAAAGAGTAACTATC |
| oligo3 | TTTCAGACAACACCAGTAATGGATCATCAGAGCTGGTCTGGTCAGTCCG |
| oligo4 | CCTAAAGCGTTAAAAGGTTTTGACGTCACTGCTTGTAGAACGTCATTTC |
| oligo5 | AGTGTTGGAACGCGTTAGTATCGCTCAAGAACGCGCCTGTTTATCAACA |
| oligo6 | GAATAGCAATATATTAATTACTGAGAATACAACATGTTCAGCGGGTAAA |
| oligo7 | AAAATCCTCTTCTGTAAGAATACGCCAACTGTCCAGACGACGTCATAAA |
| oligo8 | GATGGTGGTTTGAAATACCCAGTCGGGACGAAAATTTTGGGGGCAA |
| oligo9 | TGGAGCACTAGTGCCACGCCGGAGAGGGTAGGATAATCAATC |
| oligo10 | GCGGTCCTAATGAAACTCACAACCGAGCGTTCTTCGCGTCCGCTGC |
| oligo11 | TCAGAACTGAAAAACAGCAGGTCATTGCCTGGGTAATCAATT |
| oligo12 | CTGATTGGGGCGCCAAGCATAGTGAAATTTTTCACGGTCATAGCGC |
| oligo13 | AATCGCAAGACAAATTCCAGTTTGG |
| oligo14 | TTTTTCACCGAGATAGGGCGATGGTCAGTAAAGATTCAAAAGGAAGCCT |
| oligo15 | AATTTCACTTATAAGGGCGATATCAAACAAGGCCGGAGACAGCCCTGTA |
| oligo16 | TTTAATGGTTCCGAACCATCATCACCTTACCATCAATATGATGTTGTAC |
| oligo17 | TGTCTCACTGATCAGTTGAGGATCCCCGGGTTTAATTGGGCG |
| oligo18 | TCGACCGTGTGATAAGAGGCACCGACAA |
| oligo19 | GCCATGGTCATTTCTGCCAGCACGCCCCCTGTATTTACTCTGCTG |
| oligo20 | TATAAAGCCAACATATGCGTTA |
| oligo21 | GCCAGTACCCGCTTGGGAAGAAAGCCGGACGCCAGAGATTGT |
| oligo22 | AAGGTAAGTGTGTTCAGCAAACGCAACCAGCTTACACCGATA |
| oligo23 | CCTCAAAGAGAATACGGCATCCCGCCGCGCGCGTAATTAAAGGGCG |
| oligo24 | GGGTAGCTGTTTCCTGTAAGTGTACCAGTAAAAGAATATCAC |
| oligo25 | TCACTGTTGCCCTGCGGCCAATCCGCCGGGCGCGGT |
| oligo26 | AATTAATGCACAGTAGGGCTTAATTAGAAAAAGCCTGTAGAAAAC |
| oligo27 | GGTTTCTCATTGCAGGCGCTTTTATCAGTCGCTGAAACGAGGCATCTTT |
| oligo28 | TCGACAATAACGCCATATTTAACAAAACACCGGAATCATTTAGTT |
| oligo29 | CATCCCTGGTGTCCAGCATCAGAATTTCACGCATAAGAGGCTAAAAGAA |
| oligo30 | CTGAGTAATTCATGTAATTTAGGCAATAAGGCGTTAAAACCTAAA |
| oligo31 | AGCCGCTGCGACGAGCACGGGAGCCGCTATTGTCGGATTCTCCGTCGTT |
| oligo32 | AGTGCTCAATAATATTAAGTAGAAGATTAAGAGCCAGC |
| oligo33 | AACGCCGGTGCGTGCCTTCGAATTCGTAATCTAATGAGGCAG |
| oligo34 | GGACGCGCCTCGGGCCGTGTTATC |
| oligo35 | TCCAAAAGGAGCCTTAAAGGCCGCT |
| oligo36 | GTTGCACAGGCGGCCATCCCATCGTTAATAAAGTATTTTCGA |
| oligo37 | AGTCCGTAAAATTAAACGGGTACAATCGGCGGACATAATGCT |
| oligo38 | ACGGTACATCAAACGTAATAACCTCACCGGAGCCTCTTTAAA |
| oligo39 | GCACGCCAGCCCAGTCCGTGGAGCCGCCACGCTGGCGACGTT |
| oligo40 | GTCCTTAAATGATAGACGCCAGTGCCAAGCTGCAAGGCGAACTCACTGT |

| | |
|---|---|
| oligo41 | CCTCAGCAGCGATACCAAGCGCGAA |
| oligo42 | GCATCGGGGCTTGCAGGGAGTTTAATTGTATCGGTTCGCACTTGGT |
| oligo43 | CGGCTACACCGATATATTCGGCTTGCTTTCGAGGTGCGGGGTTTGC |
| oligo44 | CTAAAGAAACAACCATCGCCCTTAAACAGCTTGATGGCTGGATACA |
| oligo45 | AGTTTCCAAAAGCCGCGCCGACAATGACCTTTTTCCCAACCT |
| oligo46 | GACCCCCGGCACCGCTTCTGGATCTGCCAGTTTGAATCATACAACG |
| oligo47 | TACACTAAACCAGGCAAAGCGATGGGCGCATCGTAATTAGCATGCG |
| oligo48 | AAAACGACCATTCAGGCTGCGGGATAGGTCACGTTAGCCTCAATTA |
| oligo49 | GTTGTAAACGCCAGACATCACCGTTGTATGACCTGAAAACATGCAACAG |
| oligo50 | CACTCCAGCCAACGACAGTATC |
| oligo51 | ATCAACGGCGGATTGACATCAAATATTTAAA |
| oligo52 | CCCACGCCAGGGAACGGCAGCGCCATGTTTAAGTTGGGCGTG |
| oligo53 | TTAATGTGCTTTCAGAGCGGAATTTGTGAGATTCTGCTCCAG |
| oligo54 | TTTCATCCAGCTCATAGCATGAGGCTATAGGTGAGGCGGTCAAGTC |
| oligo55 | TGTTTGGGTAAACGACGTTTCTCCGTGGTGAAACA |
| oligo56 | TCTGGCCGGAACGCGATGAACAGAGTCTCACCAGCAGAAGATTAGC |
| oligo57 | AGTAGCATTAACATCCAATAAGGGGACGGCTTTCCAGCGATTAAAGACA |
| oligo58 | TTGTAAAGGGAACAGGTGCGGAAAAATACGTAATGCCACTACTGGGAAG |
| oligo59 | AATCCCCAAAAATTAATGCTGAGAGCCAGCATCGAGGTACGG |
| oligo60 | CAAGTGTAGGTTGGCAAATCAACAGACTCCAACGTCAAAGGGTTG |
| oligo61 | TTATTTCAGGCAAGGCAAAGAACCGTGCTGCCGGAAAACACTGTAGCAA |
| oligo62 | GGAGGTGAGACCTCAATCAATATCAAAACCGTCTATCAATCAAAA |
| oligo63 | ATACTTTAAATTAAGCAATAAGGTGTAGCCATTCGAAGAGGCTTGAGGA |
| oligo64 | TGATCAAATCGCTGAACCTCAAATGGCCCACTACGTGAAATCGGC |
| oligo65 | CAAAAACGAGCATAAAGCTAACGTAATGCAACTGTGAAGGCAATGAGGA |
| oligo66 | ATAAGCGATTCAACAAAAATCTAAAGCACCCAAATCAAGTTTCCTGTTT |
| oligo67 | ATGCAATGCCTGAGTAATGGATAAAAATTTTTAGAA |
| oligo68 | TTTCACCGCCTGCAACAAATCGGAACCCTATGCAGCAA |
| oligo69 | GGAAAGCGAAACCACACAGATGCCACCGTCGGTGGTGCCTTT |
| oligo70 | CTACGATTTATTTTTATAGAAAAGATTTTGTTAAAATTACAA |
| oligo71 | GGCAATGTGAGCTATCGGCCAACGCGCGGAGAGTTGGCTATTGTAT |
| oligo72 | ATTTATTTTTCCGAGTATATGTACAATTTTTGTTAAATAACA |
| oligo73 | ACCTACAGACATTCGAGCCGGAGGGTGG |
| oligo74 | GGTAAAAATCGATGAAGGGTAAAGCGGCAGCGGGTTACTGAGCCT |
| oligo75 | TTGCGTAACCAGGAGCGCGTTGCGCGTGCCAGCTGCATACGC |
| oligo76 | CTTATGTGTAGCGGTCACAGCGGTCGCAAGAATGCCAATTAA |
| oligo77 | GTAATATCATTTGCAACGCGGTCCGTTTGCGTGGTCGATCCACCGG |
| oligo78 | AACCGTCTGACACACGAAAGCCTGGCGGTTTGCGTATTCCCT |
| oligo79 | CAGAGAAGTGGAGCTTGGCCGTAATTTGCCCCAGCAGGAACC |
| oligo80 | AGAGAGGCCAGAAAAGGGAGCCCCGGGCGCT |
| oligo81 | GGCCTCTAAGGGGGAATGTGAGCGAGTACGCATTACCCGGTTCTAT |
| oligo82 | AGTATCAGAGCGTATAACGGTTGTATGCTGATTGCCGTCAGC |
| oligo83 | CCACGTGGCAAATGCGCCCGCCTGGCCCTGAGGGAGAGGGGT |
| oligo84 | GCAGTAAAACTTTTTTAACCAATATTCC |

| | |
|---|---|
| oligo85 | AACTTGCCTGCCGCCAGGATCAAATCGTCGCTGGCAGCTGCC |
| oligo86 | TGAATAATTTGCACGTAAAACAAGCGCATTAG |
| oligo87 | TCACATCAATCAGTTCAGAAAACGCGAGAGGCTTTTGCAACCCTC |
| oligo88 | ACATAAATCAGAGAAATAGCAGAAACGCAACATATAAAAGAAATATGGT |
| oligo89 | GCAGCCTATTGAGTTAAGCACGATTTTTTGTTTACCCAATCCTAAGACT |
| oligo90 | CAAAAATTGAATATGATTATCCCATAAATCAAAAACGGATGGCTAAACC |
| oligo91 | AATTTTTACACGGAATTTTTACCCTGACTATTTTGATA |
| oligo92 | TTACCTTGTGAGTGTAAACAGAAGAACGATATAGAAGGCTTAATATTGA |
| oligo93 | CATTTAATGTAAATCCTAATTGAACCTCGCCCAATAGCAAGCTAAAGGT |
| oligo94 | TCAAGAAAATTTTCCTAACGAGTTTTGACATCGTAGGAATCAACCGACT |
| oligo95 | GCAAAAGAAGAAAAACATAGCGATA |
| oligo96 | GCTAATAAACAGGGAGAAATATTGGATTATCCCCCTCAAATGGTTTTGC |
| oligo97 | TTGCTTCCAATTTCAACGGATCAAAGAAAGAAGCAAAGCGGAAGAGTAC |
| oligo98 | ATTAATTAACAAAATGAATACTAACATTAAAAAGATTAAGAGACTCCAA |
| oligo99 | ATCCTTGTGATGAAGCGCAGATATTAATGAAAGACTTCAAATATCG |
| oligo100 | ACAATGAGATAACCCACAAGATTACAGAATTT |
| oligo101 | AATAAGAACCCAAAAAATACATACATAAGACAAAA |
| oligo102 | AATTTTATCCTATTAGTTGCTA |
| oligo103 | AGTTACCAGAAGGAAACCGAGATAGCTATCTTACCATTGAGC |
| oligo104 | CCTTACGGTATTCTCCATATTATTTATCATAAATCAATATATTTTT |
| oligo105 | TTAGCAACGATTGAGGGAGGGCAGACTGTAGCGCGCATTTCGTCACCAG |
| oligo106 | TCGAGAACAAGCAAGCCGTTTATTAGAGCCAGCAAAATC |
| oligo107 | TCAATAGAAAATTCACGCAAAGACACCA |
| oligo108 | TTACCAGCCTTATTAGCGTTTCTCCCTCCGCCGCCACAAATAGTAAGCG |
| oligo109 | AAAAGGTGGCAATAATAACGGAATGCAAGAA |
| oligo110 | GGGCGACCGGCATTTTCGGTCCAGAACCCACCAGACCAGAAT |
| oligo111 | CGGAAATAATCAAGTTTGCCTTACAACGAGGAACCTTGCTCAACCGGAT |
| oligo112 | CATAAATCAGCGAGGCGTTTTAGCTGCCAGTTACAAAAAATAACC |
| oligo113 | GAATTATCACCGTAATCAGTACAGACAGTTCAGGGAGTGCCGTCAAGAG |
| oligo114 | GTCTTACCGCCCGACTTGCGGGAGGCGTCTTTCCAGAGCGTCGCT |
| oligo115 | TGAGCCACAATGAAACCATCGGTAACGACTCAGAGATAAGTAGCATAGG |
| oligo116 | GGATTATTTTAGCCTTAAATCAAGGAATCTTACCAACGCCTTAGA |
| oligo117 | CATATTCAACACGTAGAAGAACTGGCATGATAAATAAGAACCAAT |
| oligo118 | CCATTAGCAAGGCCGGAAGTCTTTCGCCACCCTGTATCATGAACGG |
| oligo119 | TACAAACTTAGCGTAAGGTAATCTTACGCAGTATG |
| oligo120 | CTGAGCCAGGATTATAAGAGGCTGAGACCCTATTTCGGAACCATAAAAC |
| oligo121 | CCCTCAGAACCCAGACGTTAGTAAA |
| oligo122 | CTTGATATTCACAAAGCATTGACAGGAGACCACCG |
| oligo123 | ATTAAAGACCACCACCAGAGCAGAGCCGCCACCCTATAGCCCCGCC |
| oligo124 | GGAAACTCAAGAGAACCCTCAGAGCCGCGCCACCCTCAGAGCTTTT |
| oligo125 | GCGGATAATAGCAAGCCCAATCCTGTAGCATTCCAGCGACAGTATT |
| oligo126 | GGTTGATCCACCACCCTCATTCCCTCATAGTTAGCATAGCAGCACC |
| oligo127 | GAATAGGTCAGAACCGCCACCTCTAAAGTTTTGTCACGTCACTTTG |
| oligo128 | TCATACAGCCTTGAGTAACAGTTCATCAGTTGAGATTTA |

| | |
|---|---|
| oligo129 | AATTTACATAAACAGTTAATGACATTATTACAGGTCTATCATAAAA |
| oligo130 | ATACATGAAAGTATGGATTAGCGGGGTTCATGTACCGTAACA |
| oligo131 | TAATCTTCGTAACAAAGCTGCTCATTATACCAGTCGCTCAACTGCT |
| oligo132 | CTGGCTGGTGAATAAGGCTTGTATGCGATTTTAAGCAACTAAGGAT |
| oligo133 | TGTACAGGAGAAACACCAGAATTTAATCATTGTGAAAGTTTCGAAG |
| oligo134 | TGAAAGAGGACAGACCGTACTCAGG |
| oligo135 | CTGAATCATTACCCTGGGAAGAAAAATCATTGCTGATTTTTGTCAG |
| oligo136 | GAACTAAAGACGACGATAATAGAGCTTATACGTTATATTATT |
| oligo137 | TTAATTTCAACCGAGTAGTAAA |
| oligo138 | GGAATTACGAGGCAGGTAATATCATTGAATACTTC |
| oligo139 | GTTTACCCGGAACACCCCCTGTCGCGCAGTCTCTG |
| oligo140 | CAGAGGGTAGTAAGAGCAACAAGAAAGATGCCCGTCGTTCCAAATCCTC |
| oligo141 | GAACTTTAAAATAATCCTGATTGTAAGAAATTGCGTAGGAGAATA |
| oligo142 | AAAATAGAGAATGAAGATGATGGCAATTGGTTTAACGTCAGAGAAAATA |
| oligo143 | AGAGGTCAATATAATGCTGTAAGGACGTAAATCAAGACAAGAGTACCAG |
| oligo144 | CCTTATAGTCACCACCAGAAGGAGTCGGGAGAAACAATATTTGAA |
| oligo145 | CTTTAATATGTTTTAAATATGAACTGGCTCATTCAACCTTCATCGAGAG |
| oligo146 | TAGTTGCATCATCATTTTGCGGAATCGCCTGATTGCTTTTAATTA |
| oligo147 | CAGGTCAAGTACGGTGTCTGGATTACCTCCCTGACACCAGGCTAGCCCG |
| oligo148 | CAAGAAGCCCTTTAAAAGTTTGAGCAAGTTACAAAATCACAAACA |
| oligo149 | TCAAAGCGAACCAGACCGATTCCATATAACAGTTGATTC |
| oligo150 | GTCATCATCATATTCCTACAGTAACAGTACCGGAAACAGTAACGT |
| oligo151 | CTTTGCCCGAACGTGGCGAATTATT |
| oligo152 | TAAAGAACGTGGTTGAAAGGAATTG |
| oligo154 | TTTGAGGATTTAGAAGTATTAGACTTTACAAACAATTCGACAACTC |
| oligo155 | TCTAAAATATCTTTAGGAGCACTAACAACTAATAGATTAGAGCCGT |
| oligo156 | AGACGCTGAGAAGAGTCAATAGTGAATTTATCAAAATCATAGGTCT |
| oligo157 | TTAACCTCCGGCTTAGGTTGGGTTATATAACTATATGTAAATGCTG |
| oligo158 | TCGGCTGTCTTTCCTTATCATTCCAAGAACGGGTATTAAACCAAGT |
| oligo159 | CCTGAACAAGAAAAATAATATCCCATCCTAATTTACGAGCATGTAG |
| oligo160 | GTATGGGATTTTGCTAAACAACTTTCAACAGTTTCAGCGGAGTGAG |
| oligo161 | ACTAAAGGAATTGCGAATAATAATTTTTTCACGTTGAAAATCTCCA |
| oligo162 | CTTAGCCGGAACGAGGCGCAGACGGTCAATCATAAGGGAACCGAAC |
| oligo163 | ACGGAGATTTGTATCATCGCCTGATAAATTGTGTCGAAATCCGCGA |
| oligo165 | GAACGAGTAGATTTAGTTTGACCATTAGATACATTTCGCAAATGGT |
| oligo166 | GCTATATTTTCATTTGGGGCGCGAGCTGAAAAGGTGGCATCAATTC |
| 25b_BioAdapt_PB | /5BiotinTEG/TTT TGCCCAGACACGTTAACCAGAG |
| 25b_ND_OvH1_PB | ACGGAGGGTAGAAGCCCTTTTTAAGAAAAGTAA TT CTCTGGTTAACGTGTCTGGGCA |
| 25b_ND_OvH2_PB | GAACCGCGCCATCTTTTCATAATCAAAATCACC TT CTCTGGTTAACGTGTCTGGGCA |
| 25b_ND_OvH3_PB | GGAGGTCAGTTGGCTTTTGATGATACAGGAGTGTAC TT CTCTGGTTAACGTGTCTGGGCA |
| 25b_ND_OvH4_PB | TAAATATGTAAAATGTTTAGACTGGATAGCGTC TT CTCTGGTTAACGTGTCTGGGCA |
| ND_Mag_Twzr_Tether1_BP | GCGCCATTGCAACAGGAAAAACTTTTCGTGCGCACCCAGGTAGACAAGGACAACT |
| Constrained Helix 0 | GAT TTA GAA GTA TTA GAC TTT ACA AAC AAT TCG ACA ACT CGT ATT GGT TAT CTA AAA TAT CTT TAG GAG CAC TAA CAA CTA ATA GAT TAG |

| | |
|---|---|
| Constrained Helix 3 | TCC GGC TTA GGT TGG GTT ATA TAA CTA TAT GTA AAT GCT GAT GCA GAT TAA GAC GCT GAG AAG AGT CAA TAG TGA ATT TAT CAA AAT CAT |
| Constrained Helix 9 | GGA ATT GCG AAT AAT AAT TTT TTC ACG TTG AAA ATC TCC AAA AAA TTT CTG TAT GGG ATT TTG CTA AAC AAC TTT CAA CAG TTT CAG CGG |
| Constrained Helix 12 | CGG AAC GAG GCG CAG ACG GTC AAT CAT AAG GGA ACC GAA CTG ACC GTA CAA CGG AGA TTT GTA TCA TCG CCT GAT AAA TTG TGT CGA AAT |
| Constrained Helix 6 | GTC TTT CCT TAT CAT TCC AAG AAC GGG TAT TAA ACC AAG TAC CGC TAA GTC CTG AAC AAG AAA AAT AAT ATC CCA TCC TAA TTT ACG AGC |
| Constrained Helix 15 | TTT TCA TTT GGG GCG CGA GCT GAA AAG GTG GCA TCA ATT CTA CTA TCT GCG AAC GAG TAG ATT TAG TTT GAC CAT TAG ATA CAT TTC GCA |

| caDNAno Helix # = Loop # | Scaffold sequence of loops (5’-3’) |
|---|---|
| **Helix 0 = loop 1** | AA TAC GAG TTG TCG AAT TGT TTG TAA AGT CTA ATA CTT CTA AAT CCT CAA ATG TAT TA TCT ATT GAC GGC TCT AAT CTA TTA GTT GTT AGT GCT CCT AAA GAT ATT TTA GAT AAC C |
| **Helix 3 = loop 2** | TG CAT CAG CAT TTA CAT ATA GTT ATA TAA CCC AAC CTA AGC CGG AGG TTA AAA AGG TA GTC TCT CAG ACC TAT GAT TTT GAT AAA TTC ACT ATT GAC TCT TCT CAG CGT CTT AAT C |
| **Helix 6 = loop 3** | GC GGT ACT TGG TTT AAT ACC CGT TCT TGG AAT GAT AAG GAA AGA CAG CCG ATT ATT GA TTG GTT TCT ACA TGC TCG TAA ATT AGG ATG GGA TAT TAT TTT TCT TGT TCA GGA CTT A |
| **Helix 9 = loop 4** | TT TTT TGG AGA TTT TCA ACG TGA AAA AAT TAT TAT TCG CAA TTC CTT TAG TTG TTC CT TTC TAT TCT CAC TCC GCT GAA ACT GTT GAA AGT TGT TTA GCA AAA TCC CAT ACA GAA A |
| **Helix 12 = loop 5** | GG TCA GTT CGG TTC CCT TAT GAT TGA CCG TCT GCG CCT CGT TCC GGC TAA GTA ACA TG GAG CAG GTC GCG GAT TTC GAC ACA ATT TAT CAG GCG ATG ATA CAA ATC TCC GTT GTA C |
| **Helix 15 = loop 6** | TA GTA GAA TTG ATG CCA CCT TTT CAG CTC GCG CCC CAA ATG AAA ATA TAG CTA AAC AG GTT ATT GAC CAT TTG CGA AAT GTA TCT AAT GGT CAA ACT AAA TCT ACT CGT TCG CAG A |

**Supplementary Table S6.** Table of the 6 loop sequences from which the force-sensing strands are designed.

| **Starting Temperature (°C)** | **Ending Temperature (°C)** | **Time spent (min) at each interval of 1°C** |
|---|---|---|
| 65 | 65 | 30 |
| 64 | 62 | 60 |
| 61 | 59 | 120 |
| 58 | 46 | 180 |
| 45 | 40 | 60 |
| 39 | 24 | 30 |
| 24 | 4 | 1 |

**Supplementary Table S7.** 2.5 day thermal annealing ramp for the base ND structure. ND samples are subjected to the following incubation cycles in order to initially disrupt any non-specific base-paring interactions and them slowly cool over ~2.5 days allowing the ND to reach its lowest energy configuration.

**Supplementary Methods.**

For the creation of our model, we split the design into two main components (see Supplementary Fig. S7). The first is the dsDNA tether, which we model as a worm like chain of contour length $L$*=1055.4nm* with persistence length $P$*=50nm*. Its force-extension behavior is thus given by

$$F = \frac{k_B T}{P}\left(\frac{1}{4}\left(1-\frac{z_{WLC}}{L}\right)^{-2} - \frac{1}{4} + \frac{z_{WLC}}{L}\right) \quad (1)$$

where $k_B$ is the Boltzmann constant, and $T$ is the absolute temperature.

The other component is the NanoDyn on the magnetic bead. We assume that the magnetic bead is held fixed by the magnetic field and model the NanoDyn as a stiff rod of length $l$. We do not know on which point of the bead the attachment between the bead and the NanoDyn is located and denote the unknown angle between the horizontal and the line from the center of the bead to the attachment point of the NanoDyn as $\Theta_0$. We then assume that the NanoDyn is oriented radially away from the bead in the absence of a force but that the attachment between the NanoDyn and magnetic bead has some ability to bend. We thus model the connection between the NanoDyn and the bead as a torsional spring with spring constant $K_\theta$ while we denote the actual angle between the horizontal and the axis of the NanoDyn as $\Theta$. In order to avoid numerical instabilities stemming from the fact that a fixed NanoDyn length $l$ results in a maximum extension of the entire system reached for $\Theta = 90°$, we artificially allow the NanoDyn length $l$ to vary by treating the NanoDyn extension as a very stiff spring with rest length $l_0$*=100nm* and spring constant $K_l$*=10000pN/nm.* This in principle allows arbitrary extensions while still keeping the actual length $l$ very close to its equilibrium value $l_0$ for any reasonable force. With all these parameters, the extension of the NanoDyn in the vertical direction is

$$z_{tor} = l\sin(\theta) - l_0 \sin(\theta_0) \quad (2)$$

and the potential energy of this system is given by

$$V(l,\theta) = \frac{K_l}{2}(l-l_0)^2 + \frac{K_\theta}{2}(\theta-\theta_0)^2 - F \cdot z_{tor}.$$

In equilibrium, this potential energy will be minimized with respect to $l$ and $\varphi$ and we can thus take the derivative with respect to these two dynamic degrees of freedom separately to yield

$$F = \frac{K_l(l-l_0)}{\sin(\theta)} \quad (3)$$

$$F = \frac{K_\theta(\theta-\theta_0)}{l\cos(\theta)} \quad (4).$$

We can lastly represent the full length of the system by the sum of its components:

$$z = z_{tor} + z_{WLC} + z_0, \quad (5)$$

where $z_0$ is an unknown offset required since the instrument can only measure relative distances. This gives us five equations Eqs. (1-5) for the five unknowns $z_{tor}, z_{WLC}, l, F$,and $\Theta$. We can simplify this system of equations somewhat by eliminating the force between equations (1), (3), and (4) to leave us with four equations and unknowns. This set of four equations allows us to numerically solve for the four unknowns using the python function optimize.root for any experimentally measured $z$ and any given set of the parameters $z_0$, $\Theta_0$, and $K_\theta$, where we restrict the possible angles $\Theta$ for the torsional spring to the interval between -π/2 and π/2. Once the four unknowns are determined, we calculate the force $F$ based on Eq. (3) or (4) (choosing Eq. (4) for values of Θ far away from zero and Eq. (5) for values of $\Theta$ close to zero to maintain numerical stability). This numerically yields a force-extension curve *F(z)* for a given set of parameters $z_0$, $\Theta_0$, and $K_\theta$. Finally, we use least squares fitting (python function curve_fit) to find the parameters $z_0$, $\Theta_0$, and $K_\theta$ that minimizes the difference between the experimental data and the numerically calculated force extension curve.

To then fit this model to force extension curves featuring a rupture event, additional steps are needed. We first need to find the size of the rupture and the rupture force. We calculate the rupture force as described above. We then take the differences between each consecutive extensions $z$ and use our previously found rupture force to find the region just as the rupture begins (the first data point where the force exceeds the rupture force) and record the points corresponding to a set of at least 20 consecutive differences larger than 20nm and iterating on this until the following difference goes below the 1nm threshold. We can then use this set of large differences and subtract its sum (the total difference in extension across the entire rupture region) from the high force regime data. For all data points in the rupture region itself the extension is set to the extension at the beginning of the rupture region. Then, a force extension curve is fitted as described above to the data, from which the rupture has been removed. In the absence of a detailed understanding of the mechanics of the flexible part of the open NanoDyn, we finally reintroduce the rupture by extending the tether for extensions beyond the rupture point. As we know that only a fraction of the total length of the tether is contributing to the extension of the system at a given finite force, we must again numerically solve the force-extension equation of the worm like chain

$$F_{rup} = \frac{k_b T}{P}\left(\frac{1}{4}\left(1-\frac{\Delta z}{\Delta L}\right)^{-2} - \frac{1}{4} + \frac{\Delta z}{\Delta L}\right)$$

to obtain the extension change $\Delta L$ of the worm like chain required to result in a $\Delta z$ jump of the measured extension at the rupture force $F_{rup}$.
